# Supercharged binding modules can modulate engineered poly(ethylene terephthalate) hydrolase thermostability and functional persistence

**DOI:** 10.64898/2026.05.24.727315

**Authors:** Antonio DeChellis, Sumay Trivedi, Lingjun Xie, Sagar Khare, Shishir P. S. Chundawat

**Author notes:** **Corresponding Author** - Shishir P.S. Chundawat.

## Abstract

Poly(ethylene terephthalate) (PET) is a highly recalcitrant polyester plastic whose resistance to degradation has contributed to widespread environmental accumulation. Enzymatic PET depolymerization has emerged as a promising bioremediation strategy, but PET hydrolysis remains challenging due to the insoluble and semi-crystalline nature of PET and the poor thermostability of many PET hydrolases at elevated temperatures. Here, several electrostatically supercharged PET binding modules (PBM) were fused to a PET-hydrolyzing Cutinase Catalytic Domain (CD) from the thermophilic microbe *Thermobifida fusca* to investigate how engineered PBM surface charge influences PET hydrolysis behavior. All PBM designs were derived from a native *T. fusca* family-2a carbohydrate binding module (CBM) as starting template. Since PET exhibited a substantially negative zeta potential, and accordingly, all positively supercharged PBMs displayed the strongest PET binding interactions in pull-down binding assays. However, stronger PET binding did not translate to improved hydrolysis activity for the fusion constructs. Instead, a slightly negatively charged PBM-CD fusion (D2 construct) exhibited activity comparable to the Cutinase CD on finely milled PET powder while showing substantially improved activity on intact PET discs, suggesting potential advantages for depolymerization of minimally processed PET feedstocks. Thermostability analysis identified an approximately 10 °C increase in melting temperature for the D2 fusion construct, corresponding to enhanced catalytic persistence and a shifted optimal hydrolysis temperature. Consequently, this construct exhibited an approximately 2-fold increase in long-term hydrolysis activity on milled PET and up to a 10-fold increase on intact PET discs, even at high solids loadings, compared to the native Cutinase CD. Collectively, these findings demonstrate that thermostability, rather than adsorption to PET alone, is a dominant factor governing functional persistence of PET hydrolases.

## INTRODUCTION

Overproduction of single use plastics has culminated into a global crisis as billions of tons of plastic waste enters ecosystems yearly, posing environmental and human health risks each year.^1–3^ Plastic production has exploded in the 21^st^ century with consistent demand increasing, leading to much of this single use plastic accumulating environmentally.^2,4^ One such plastic, poly(ethylene terephthalate) (PET), is a major contributor to this issue as one of the most widely produced single-use plastics.^2,5^ PET is a petroleum derived polyester thermoplastic prized for its stable semi-crystalline structure, high tensile strength, and inherent chemical resistance commonly employed for single use packaging for food and beverage, as well as for polyester textile fibers.^4,6^ These properties that brand PET as an attractive industrial plastic also result in remarkably slow decomposition rates, causing PET to accumulate as a persistent environmental contaminant.^4^ One potential avenue for environmental remediation is the enzymatic depolymerization of PET using esterase PET hydrolases capable of cleaving ester linkages in the PET backbone.^7,8^ This solution has gained particular interest in the field after Yoshida et al. 2016^8^ isolated a novel microbe, *Ideonella sakaiensis*, that degrades PET for assimilation of carbon into its metabolism. This microbe secretes the PETase and MHETase enzymes that work in conjunction to depolymerize PET into its constituent terephthalic acid (TPA) and ethylene glycol monomers.^8^ These enzymes are quite limited, however, as they possess poor thermostability preventing them from effectively processing PET at higher temperatures where it is more amenable to enzymatic access.^7^Although enzymatic PET depolymerization is a promising strategy for closed-loop recycling, there are still several challenges preventing widescale implementation.

Efficient PET hydrolysis presents unique challenges compared to soluble substrates due to the insoluble and semi-crystalline nature of PET. PET is highly insoluble, thus productive hydrolysis with a soluble enzyme depends on adsorption and persistent interaction with the PET surface.^9^ This hydrolysis efficiency generally increases at temperatures approaching the polymer glass transition temperature (*T_g_* = 80 °C) where water plasticizes polymer chains, increasing polymer chain mobility, promoting improved substrate access.^7,10,11^ However, novel PETase rapidly loses activity at these elevated temperatures due to the aforementioned poor thermostability.^8^ As an alternative, researchers have sought other potential PET hydrolases from thermophilic microbes to identify esterase targets with more robust temperature resistance. Cutinase enzymes, identified in both bacteria and fungi, are one such target that naturally catalyze ester bonds within cutin, the waxy biopolyester found naturally occurring in the cuticle of plants.^12,13^ Several Cutinase enzymes with superior thermostability than PETase have shown modest PET hydrolase activity at temperatures exceeding 50 °C, including a Cutinase from the thermophilic microbe *Thermobifida fusca* as well as Leaf-branch compost Cutinase (LCC).^14–17^ Although a more stable starting point for PET hydrolase candidates, these enzymes still suffer from the same limitations: i) adsorption challenges to the insoluble PET substrate, and ii) thermal persistence at the high temperatures needed for effective PET hydrolysis.

Much effort has been placed on improving both *Ideonella* PETase as well as various Cutinase enzymes to overcome their limitations. Computational redesign has been a successful tool to stabilize PETase, with several thermostable variants having exhibited vast improvement from the native PETase construct.^18–21^ Similar efforts have been applied to LCC, with the field-leading LCC ICCG variant having resulted in melting temperatures beyond the PET glass-transition temperature and up to 90% conversion of PET.^16^ In order to address limitations related to interfacial catalysis, much of the field has sought to append cellulose binding carbohydrate binding modules (CBMs) to PET hydrolases in order to improve substrate targeting drawing similarities between the molecular backbone of PET and cellulose.^22^ Type-A CBMs in particular possess surface exposed planar aromatic residues known to align to planar glucan rings within cellulose, enabling binding through π-π ring stacking interactions.^23,24^ These same interactions have been shown to specifically bind to PET through the same mechanism, validating the application of this engineering method.^25,26^ Previous works focused on appending different CBMs to various PET hydrolases have shown modest improvements in hydrolytic activity at low solid loadings, but the benefit of the added CBM has shown to be minimized at high solid loading concentrations.^27–29^ Thus, more work needs to be done in order to design CBMs that can target PET effectively enough for industrially relevant conditions.

One potential method of making CBMs more amenable for PET recognition, and engineer novel PET Plastic Binding Modules (PBMs), is through electrostatic surface engineering, or protein supercharging. This rational design approach involves mutating several different solvent exposed amino acid residues to either negative (D,E) or positive (R,K) residues to generate protein surfaces with high theoretical net charge.^30^ We have previously applied this method to a family-2a CBM from the thermophilic microbe *T. fusca* naturally appended to a cellulase degrading enzyme Cel5A.^31^ As a result of supercharging, we observed a significant improvement in cellulose binding affinity for positively charged CBMs, improved thermostability in the presence of substrate, as well as modulation in pH optima, ultimately resulting in significant increases in cellulose hydrolysis yields.^31^ These observations were conceptualized within the framework of the Sabatier principle^32^, where intermediate-strength substrate interactions maximize catalytic turnover. This principle has been applied to several relevant interfacial enzymes to dictate heterogenous catalysis behavior^33,34^, and within the context of charge engineering, a critical net charge can be identified where catalysis is maximized.

Here, we build upon our previous work to implicate charge engineered CBMs as novel PBMs for enabling PET depolymerization. *We hypothesized that supercharging could be used to tune CBM electrostatic surface properties to maximize PET association, resulting in improved PET hydrolysis activity.* To investigate this hypothesis, a series of charge-engineered *T. fusca* CBM2a variants from our previous work spanning a broad net charge range were fused to the catalytic domain of *T. fusca* Cutinase and systematically characterized. PET hydrolysis activity was assessed across multiple PET substrates and reaction conditions, while differential scanning fluorimetry, thermal inactivation assays, and GFP-based binding module pull-down binding studies were used to evaluate the relative contributions of substrate interaction and enzyme stability to hydrolytic performance. These studies provide insight into how electrostatic engineering of appended binding domains influences PET depolymerization behavior in heterogeneous interfacial biocatalysis systems.

## EXPERIMENTAL SECTION

### Reagents

The polyester terephthalate (PET) plastic used in this study was procured from Goodfellow as 0.25 mm thick amorphous films (252-144-75) and semi-crystalline (>50%) 300 µm powder. PET films were washed with 1% sodium dodecyl sulfate (SDS) and ample deionized (DI) water and allowed to dry completely before being punched into discs using a 1/8” circle hole punch (CARL) for usage in assays. Powder substrate contained a significant amount of PET fines that were unable to settle via centrifugation. Thus, the PET powder was sieved using a No. 100 (150 µm) fine mesh sieve (McMaster – Carr) to remove fines, and the powder was washed to remove manufacturing contaminates through five successive wash steps of vortexing with 50 mL of 1% SDS, centrifugation to settle solids, and decanting the supernatant. SDS was effectively removed by ten successive washes with 50 mL DI water. The final wash step was filtered through Whatman Grade 1 filter paper using a Buchner funnel until the final washed and sieved powder was completely dry. For usage in assays, PET powder was resuspended in DI water to make a 100 g/L slurry. Genes containing the CBMs used in this study were originally synthesized via a Department of Energy Joint Genome Institute (DOE-JGI) gene synthesis grant from our previous work.^31^ Genes for the wild-type *Thermobifida fusca* Cutinase catalytic domain (CD) were synthesized and cloned into a pET-26b(+) expression vector (https://www.addgene.org/vector-database/2563/) by Twist Bioscience. All other reagents used were procured from either Sigma Aldrich or Fisher Scientific unless otherwise noted in subsequent sections.

### CBM-Cutinase Chimera Library Construction

CBM2a-Cutinase chimeras were generated via sequence and ligation independent cloning (SLIC)^35,36^ by inserting the Cutinase CD into vectors containing the native and supercharged CBMs, along with their linkers. The process for rational computational design of the negatively (D1, D2) and positively (D3, D4) supercharged CBMs is detailed in our previous works.^31,37^ The cloning job discussed herein involved removing a Cel5A carbohydrate active enzyme catalytic unit from the original pET-45b(+) vectors from previous work^31^ to insert the Cutinase subunit into each vector on the C-terminus. Briefly, plasmids were extracted and purified for the original WT and D1-D4 – Cel5A constructs, as well as the Cutinase plasmid using miniprep kits (IBI Scientific). Individual backbone linearization primers were designed to bind directly downstream of the C-terminal stop codon in the forward direction, and at the end of the linker peptide covering the CBM in the reverse direction with overhangs complimentary to the Cutinase insert. Insert primers were designed to cover the Cutinase CD with forward insert primers containing an overhang complimentary to the CBM2a linker, and reverse primers containing an overhang complimentary to the pET-45b(+) vector of the final chimera. Although all five final constructs contain the same linker peptide found native in the CBM2a – Cel5A enzyme, codon optimization of each original individual CBM construct prevented the creation of universal insert primers. All primers were designed to possess a consistent 65°C melting temperature for standardization of annealing settings for PCR.

Polymerase chain reactions (PCR) were run for all five CBM and linker backbones as well as five different iterations for the Cutinase insert to generate the proper overhangs for each cloning job. Vector and insert PCR was setup using 5 µL each of 5 mM forward/reverse primer prepared in DI water, 100 ng of template DNA, 25 µL 2x Phusion Master Mix (New England Biolabs), and the volume was made up to 50 µL with PCR grade water (IBI Scientific). Following an initial denaturation at 98 °C, reaction mixtures were subject to 35 cycles consisting of: i) denaturation at 98 °C for 10 s, ii) annealing at 60 °C (T_m_ – 5°C) for 30s, iii) elongation at 72 °C for 60s/kbp and a final elongation of 5 min for inserts and 10 min for vectors in an Eppendorf 6377 Mastercycler Nexus. PCR product was purified using a PCR cleanup kit (IBI Scientific) following manufacturer protocols and product concentration was measured using a SpectraMax M5e spectrophotometer with true concentration identified by correcting for contamination based on 230 nm absorbance.

Prior to conducting the SLIC reactions, template DNA was first degraded via Dpn1 digestion by combining 100 ng vector and insert products in a 2.5, 5, and 10 I:V product molar ratio with 1 µL of 10x Cut Smart Buffer (New England Biolabs, NEB) and 1µL of Dpn1 enzyme (NEB) with the final volume made up to 10 µL with PCR water. DPN1 digestion was conducted at 37 °C for 2 hours then placed on ice. Once cooled, 0.5 µL of T4 DNA polymerase was added to each tube along with 2 µL of NEB 2.1 buffer and volume made up to 20 µL total with PCR water. Tubes were incubated for 5 min at 25 °C then immediately placed on ice before transforming the entire SLIC mixture into E. cloni competent *E. coli* cells and plating cells on LB-agar plates supplemented with 100 µg/mL of carbenicillin selection antibiotic. Plates were incubated at 37 °C for 16 hours, and colony PCR was used to identify colonies containing the desired plasmids. Plasmids from colonies producing verified sequences (Sanger sequencing, Azenta) were extracted and transformed into *E. coli* BL21 (DE3) competent cells (NEB) for expression.

### Recombinant Protein Expression and Purification

The native (WT) CBM2a – Cutinase fusion, all four supercharged variant fusions (D1-D4 CBM2a – Cutinase) and the isolated *Tf.* Cutinase CD constructs were all expressed in *E. coli* BL21 (DE3) cells. Glycerol stocks for each construct were used to inoculate 10 mL of standard LB media supplemented with 100 µg/mL of carbenicillin for CBM fusion constructs (pET-45b) and 50 µg/mL kanamycin for the isolated cutinase CD (pET-26b). Starter cultures were incubated overnight for 16 hr at 37 °C and 200 RPM orbital mixing in an Eppendorf Innova incubator. After overnight growth, the entire inoculum was transferred to 500 mL sterile LB media with the appropriate antibiotic per construct and incubated again at 37 °C for 2 – 4 hr until the cultures reached mid-log phase growth (0.4 – 0.6 OD_600_). Once cultures reached mid-log phase, cultures induced with stock 1 M Isopropyl β-d-1-thiogalactopyranoside (IPTG) such that the effective concentration of inducer in the culture was 0.4 mM and incubated for an additional 24 hr at 16 °C with 200 RPM orbital mixing. After expression, cells were isolated from spent media via centrifugation at 30,100 x g for 10 mins at 4 °C. Isolated cell pellets were resuspended by vortexing in 15 mL cell lysis buffer (20 mM phosphate buffer, 500 mM NaCl, and 20% (v/v) glycerol, pH 7.4), 200 µL protease inhibitor cocktail (1 µM E-64, Sigma Aldrich), and 15 µL lysozyme (Sigma-Aldrich) for every 3 grams of dry cell pellet. Lysis was done on ice using a Qsonica Q700 sonicator fit with a 1/4” microtip probe for 5 minutes total runtime (Amplitude = 10, pulse on time: 10s, pulse off time: 30s). Soluble cell lysates were isolated from insoluble cell debris via centrifugation at 12,000 x g for 45 min at 4 °C. Lysates were decanted from the pelleted cellular debris and clarified via filtration with a 0.45 µm syringe filter before downstream purification.

Immobilized metal affinity chromatography (IMAC) was utilized to isolate N-terminal HIS-tagged CBM-Cutinase fusions or the C-terminal HIS-tagged Cutinase catalytic domain from genomic *E. coli* proteins. Briefly, IMAC was performed on a Bio-Rad NGC fast protein liquid chromatography (FPLC) instrument fit with a His-trap FF Ni^2+^ – NTA column (Cytiva). Prior to sample application, the column and system plumbing were equilibrated with 5 column volumes (25 mL) of start buffer A (100 mM MOPS, 500 mM NaCl, 10 mM imidazole, pH 7.4) at 5 mL/min. Elution buffer B (100 mM MOPS, 500 mM NaCl, 500 mM imidazole, pH 7.4) was added directly to cell lysates so that the effective concentration of imidazole in the samples for column binding was 20 mM. Samples were loaded onto the column at a constant rate of 2.5 mL/min, and once the full sample was applied, start buffer A was flushed through the column at 5 mL/min to elute nonspecifically bound proteins off until a stable baseline was achieved according to the A_280_ chromatogram. Elution was performed in two isocratic steps: i) with 5% elution buffer B at 5 mL/min and ii) 100% elution buffer B at 5 mL/min, with fractions collected based on A_280_ peak. Identity and purity of final collected fractions was confirmed via SDS-PAGE. Samples with >90% purity were desalted into 50 mM HEPES pH 8.0 + 100 mM NaCl using PD-10 desalting columns (Cytiva) and concentrated to > 3 mg/mL in a Vivaspin 20 10 kDa cutoff centrifugal concentrator (Cytiva) before long term storage at -80 °C.

### PET Degradation Assays and Product Detection

Hydrolysis of PET substrates and the subsequent determination of the terephthalic acid (TPA) byproduct were used to assess the activity of the enzymes used in this study. Hydrolysis of amorphous PET discs and semi-crystalline PET slurries were performed in 0.2 mL round bottom microplates (Greiner BioOne) with a constant 4 mg of PET used for both substrates. The base hydrolysis scheme was formulated as follows. Plates were prepared by adding either 40 µL of 100 g/L PET slurry or by individually placing two-hole punched discs in each well using tweezers with 35 µL of DI water. Enzymes were diluted in buffer to a reach a constant 120 nmol / g PET loading in each well and 50 µL of dilution was added to PET containing wells. Assays sweeping pH utilized 50 mM buffers ranging from pH 5.0 – pH 7.5, with all assays afterwards sticking with a standard 50 mM phosphate pH 6.0 buffer based on results discussed in the subsequent sections. Reaction volume was made up to 200 µL with DI water, plates were sealed with TPE capmat-96 (Micronic) plate seals and taped tightly with packing tape on all edges to prevent evaporation. Unless noted otherwise elsewhere, plates were incubated for 24 hr at 50 °C based on the published temperature optimum of *Tf.* Cutinase^15,38^ with 5 RPM end-over-end mixing in a VWR hybridization oven. Variations of this base hydrolysis schematic included increasing incubation temperature, time, and substrate loading, with all other variables held constant. In all cases, each condition was tested in quadruplicates.

TPA product was detected via plate based fluorometric assays by converting TPA to the fluorophore hydroxyphthalic acid (HOTP) using a modified Fenton reaction. This detection assay was modified from previously published^39,40^ protocols to improve low end sensitivity and substantially increase the linear detection range. Reaction plates were stored at -20 °C after hydrolysis to arrest the enzymatic reaction before TPA detection. Plates were thawed and centrifuged at 3,000 x g to settle all PET solids before aliquoting 100 µL of hydrolysate to 0.2 µL PCR tubes. Hydrolysate aliquots were heated at 95 °C for 10 mins, then cooled to 10 °C for 20 mins in a thermocycler in order to denature enzyme remaining the hydrolysates. PCR tubes were spun down in a microcentrifuge to pellet denatured proteins and 90 µL of the clarified supernatant was added to Fluotrac 96-well flat bottom black microplates (Greiner Bio-One). The following reagents were added in this specific order: i) 10 µL of 1 M NaOH for pH adjustment, ii) 40 µL 3.5% H_2_O_2_, iii) 30 µL 5 mM EDTA, and iv) 30 µL 5 mM FeSO_4_. The addition of hydroxide is necessary for pH adjustment after the hydrolysis assay, and the addition of peroxide is necessary for boosting sensitivity by improving hydroxy free radical formation. As FeSO_4_ is susceptible to oxidation in water, fresh dilutions were prepared from stock reagent each assay in acidulated DI water. Upon addition of all reagents, plates were sealed with aluminum plate seals and incubated 30 min at 25 °C. After incubation, fluorescence of the HOTP product was measured in a SpectraMax M5e spectrophotometer (Ex = 320 nm, Em = 430, Cutoff = 420 nm). Measured fluorescence was compared to TPA standards at known concentrations subjected to the same sample handling as hydrolysates.

### Thermal Shift Assays and Data Analysis

Melting temperatures for all five fusion constructs and the isolated Cutinase CD was measured via differential scanning fluorimetry (DSF) in a Bio-Rad CFX Opus 96 Real-time PCR system following previous protocols.^37^ Briefly, 17.5 µL of DI water, 25 µL of 10 µM protein dilution, and 2.5 µL of 1 M buffer ranging from pH 5.0 – 7.5 were added to a clear low-profile Hard-Shell 96-well PCR plate (Bio-Rad). Once mixed, 5 µL of 50x SYPRO orange dye (Thermo Fisher Scientific) was added and plates sealed with a clear adhesive Microseal ‘B’ plate peal (Bio-Rad). Protein melt curves were generated by heating samples from 25 – 99 °C at an increment of 0.5 °C with 5 s read time at each temperature. Each enzyme was tested in quadruplicates with buffer only blanks prepared for each pH tested. Raw data was exported into MATLAB and interpolated using MATLAB’s interp1 function with a piecewise cubic hermite interpolating polynomial to interpolate data between 0.0001 °C. To identify melt transitions, MATLAB’s gradient function was used to evaluate the numerical derivative of the raw data. The raw derivative values were multiplied by a scalar of -1 to compute the negative derivative and interpolated in the same manner described earlier. Melting temperatures were identified using MATLAB’s min function to identify peaks in the negative derivative curves.

### Thermal Inactivation PET Assays

To corroborate DSF findings and assess if measured melting temperature improvements are consistently observed in enzyme function, the native CD and thermostabilized D2 CBM2a – Cutinase fusion construct were preincubated at different temperatures before hydrolyzing PET. Enzyme dilutions were prepared in 50 mM phosphate, pH 6.0 and 60 µL of dilution were placed into PCR tubes. Tubes were heated at temperatures ranging from 45 – 80 °C for 30 min in a thermocycler in quadruplicate, with control aliquots beings incubated at room temperature for the same time scale. After this pre-incubation, enzyme aliquots were centrifuged to pellet any denatured protein and the clarified supernatant immediately transferred to microplates to assay residual activity of soluble enzyme in the same manner described as above. Data was fit to a four-parameter logistic curve to identify the inflection point corresponding to the temperature where the enzyme loses 50% of its normal activity (T_50_).

### Soluble Substrate Kinetic Assays

Kinetic curves were constructed using 4-nitrophenyl acetate (pNPA) as substrate for the native CD and improved D2 CBM2a – Cutinase fusion construct to assess potential changes in the catalytic rate or mechanism. Stock pNPA solutions were prepared in pure DMSO at concentrations of 50, 10, and 2.5 mM respectively in order to test substrate concentrations ranging from 5 – 750 µM. To conduct kinetic assays, an appropriate amount of substrate was added to 37 µL DI water and 10 µL 1 M phosphate pH 7.0 in clear flat bottom microplates. A pH of 7.0 was utilized in order to visualize real time formation of pNP product without needing to quench with an alkaline solution. Additional DMSO was added to wells as needed in order to maintain a consistent amount of the solvent across all wells. Enzyme concentration was fixed at 0.05 nmol per well and prepared as 150 µL dilutions in DI water. Enzyme dilutions were added last before immediately reading absorbance at 410 nm for 10 mins with a 10 s interval, with a carriage mixing step include between measurements inside a SpectraMax M5e spectrophotometer. Assays with pNPA were extremely sensitive, since the substrate will auto-hydrolyze in the presence of water. Thus, a low enough concentration was used to measure the linear range of the progression curve. Due to substrate instability, a substrate concentration above 750 µM could not be reliably tested.

### GFP Pull-down Binding Assays

N-terminal CBM tagged GFP fusions were constructed for each CBM used in this study from our previous work^31^ and used to assess binding to PET substrates. The most negative D1 CBM2a fusion was omitted due to challenges related to insoluble expression in *E. coli*. Due to substantial non-specific absorption of isolated GFP, as well as poor binding with bovine serum albumin (BSA) present, full-scale binding isotherms were not constructed. Instead, binding was measured at one fixed concentration, (1 µM) both with 0.05% (w/v) BSA and without BSA present. At higher concentrations, the signal to noise ratio exponentially increased, limiting the effective concentration range where binding can be assessed. Assays were conducted in 0.2 mL round bottom microplates (Greiner Bio-one) with 40 µL of 100 g/L PET slurry or 2 PET with 35 µL of DI water in binding wells for a total load of 4 mg PET. Buffer concentration in each well was fixed with 20 µL of 0.5 M HEPES pH 6.5 stock based on preliminary binding screens. Protein dilutions were made in water at a concentration of 1.43 µM such that the addition of 140 µL of dilution to each well would bring the final reaction volume to 200 µL, supplying an effective protein concentration of 1µM. Substrate blanks replaced PET substrates with 40 µL DI water, with all other reagent additions held constant. These blanks were used to measure baseline GFP fluorescence for each construct without PET present. For assays including 0.05 % (w/v) BSA, 20 µL of stock buffer containing 0.5 M HEPES pH 6.5 + 5 mg/mL BSA was used, with all other regent volumes kept constant. All conditions were tested with six replicates, for both binding wells containing PET substrates, and standard wells for each enzyme without PET. Plates were sealed with a TPE capmat-96 (Micronic) plate seal and incubated at 25 °C with 5 RPM end-over-end mixing in a VWR hybridization oven for 15 mins. After incubation, plates were centrifuged at 3,000 x g to settle PET substrate and 100 µL of supernatant was transferred to Fluotrac 96-well flat bottom black microplates (Greiner Bio-One) to measure GFP fluorescence. Binding was quantified by assessing the depletion of the GFP signal from PET containing wells compared to substrate blank wells. After the this initial binding assessment, 100 µL of fresh buffer (100 mM HEPES pH 6.5 or 100 mM HEPES + 1 mg/mL BSA) were added to each well (binding wells and substrate blank wells), to supplement the 100 µL and plates were allowed to re-equilibrate by incubating at 25 °C with 5 RPM end-over-end mixing for 15 mins once more. Net recovery of GFP fluorescence was used to report outcomes from re-equilibration calculated as the difference in GFP fluorescence between re-equilibration and initial binding supernatants (accounting for dilution), normalized to the initial substrate blank fluorescence. Based on this calculation, a positive net recovery indicates desorption of bound protein from PET, and negative net recovery indicating further adsorption to PET during re-equilibration.

## RESULTS AND DISCUSSION

### Supercharging Does Not Meaningfully Alter pH-Dependent PET Hydrolysis

It was initially hypothesized that generating positively charged binding surfaces would ultimately produce chimeras with improved hydrolysis activity. Previous work with these same CBMs fused to a cellulase showed that positive supercharging was successful for this exact reason; imparting positively charged residues bolstered higher binding affinity to substrate which increased hydrolysis yields, shifted pH optima, and stabilized the enzyme-substrate complex at higher temperatures.^31^ This enhancement in activity follows the Sabatier principle^32^, where a critical, substrate specific enzyme net charge produces an intermediate binding strength where neither adsorption nor desorption are limiting, thus maximizing enzymatic turnover. Many similarities have been drawn between PET and cellulose, specifically regarding the presence of planar aromatic rings on a molecular level^22^ that Type -A CBMs like those used in this work have been shown to specifically bind to.^25^ This specific binding is highly temperature and substrate dependent, where binding is improved at elevated temperatures where polymer chains may be more accessible and the PET surface becomes more crystalline.^41^ Based on zeta potential measurements (**Supplemental Figure S1**), both substrates share a similar electrostatic profile as well, with PET possessing a much more negative surface as compared to cellulose. For these reasons, it was initially hypothesized that positively charged CBMs, D3 and D4, would likely be the best fusion partners for increased PET binding and hydrolysis, if adsorption was rate-limiting.

Results in **Figure 2** directly conflict with this hypothesis. On milled, semi-crystalline PET (**Figure 2A**), the Cutinase CD shows the highest catalytic performance, with only the slightly negative D2 CBM2a fusion being nearly comparable, with slightly higher activity at pH 6.5 as well. More importantly, there is no shift in the pH optima behavior, instead, the optimum of the catalytic domain dictates the optimum of the full-length enzyme. Previous binding assays on cellulose have shown that specific binding for these particular CBMs is best around a more acidic pH of 5.0 – 5.5.^31^ Therefore, it is possible that this mismatch in pH dependent behaviors, binding and catalysis, may be limiting improvements in productive binding to the PET surface. While the CBM is likely contributing to increased adsorption to PET, data suggests that it is likely doing so in a nonproductive manner where either only the CBM is binding, or the full-length enzyme is adsorbing in a manner where the Cutinase domain is in a non-catalytically relevant conformation. This could explain the significant decrease in activity observed for all fusions as compared to the CD alone. Milled PET provides a large accessible surface area where nonproductive adsorption by a CBM could decrease hydrolysis yield. Conversely, with high surface area, binding could become less of a limiting factor for the isolated CD, producing the high hydrolysis yield observed.

**Figure 1.**
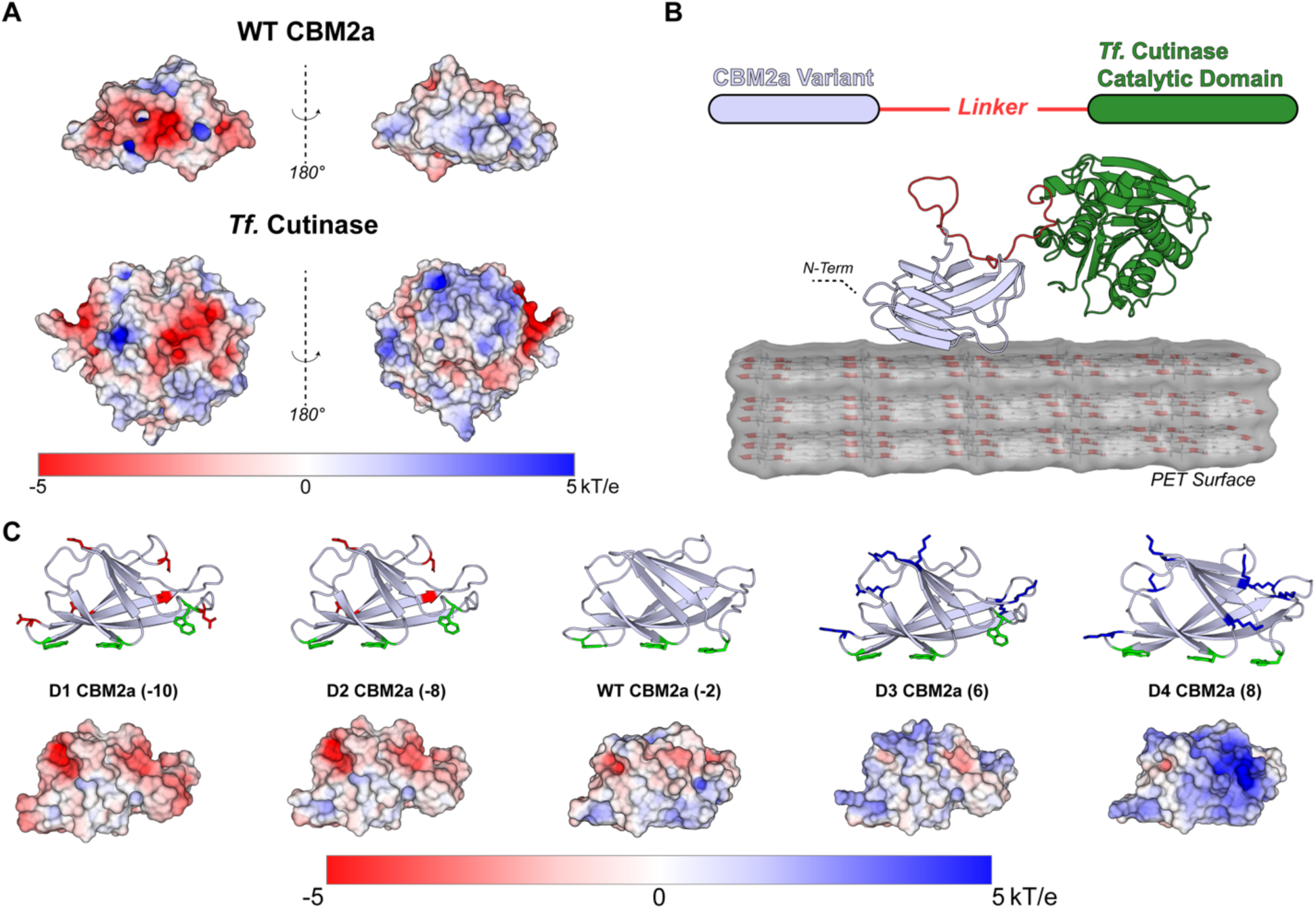
Library design and electrostatic characterization of charge-engineered CBM2a – Cutinase fusion constructs. (A) Electrostatic surface (ES) maps of the native *T. fusca* family – 2a binding module (CBM2a) and *T. fusca* Cutinase catalytic domain (CD). Protein surface models are oriented such that the planar CBM binding face is oriented downward, and the Cutinase active site is oriented towards the viewer, with alternative views provided by rotating 180° about the y-axis. Protein structures were generated with Alpha Fold, and output structures were relaxed in Rosetta. ES maps were generated in PyMol using the Adaptive Poisson – Boltzmann Solver (APBS) plugin with surface maps ranging from -5 kT/e (red) to +5 kT/e (blue). (B) Schematic representation of the engineered CBM2a-Cutinase fusion enzyme designs used in this study. N-terminal charge-engineered CBM2a variants were fused to the *T. fusca* Cutinase catalytic domain through the same flexible linker found native in the original CBM2a – Cel5A enzyme that the CBM2a used in this work originates from. A conceptual model of PET surface engaged with the fusion enzyme construct is shown here. The full-length protein model was generated in Alpha Fold and relaxed using both the AMBER forcefield and Rosetta. (C) Visualization of the CBM2a supercharged variants, and native CBM2a protein used in this study. Protein structures were generated in Rosetta during the computational design process. Cartoon models highlight planar aromatic binding residues (green) and as well as mutations shown to negative (red) or positive (blue) residues. ES maps have been provided for each enzyme and are oriented in the same pose as their corresponding cartoon structure. All structures were visualized in PyMOL.

**Figure 2.**
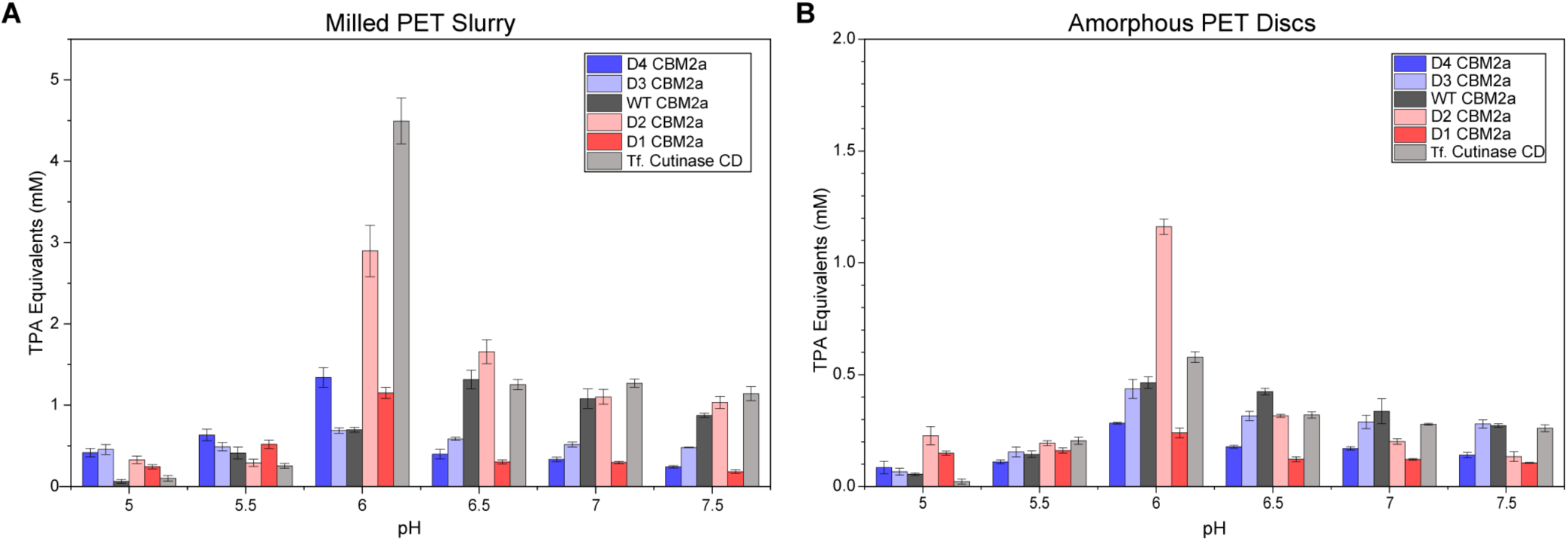
Supercharged CBM2a – Cutinase fusions do not display shift in optimal hydrolysis pH. Hydrolysis of (A) semi-crystalline milled PET and (B) amorphous PET discs at different pH reaction conditions. Substrate amount was fixed at 4 mg using milled PET prepared as a slurry or amorphous sheets hole punched into 3 mm discs. All enzyme dilutions were made in DI water and a constant enzyme-substrate loading of 120 nmol / g substrate was used for assaying activity. Hydrolysis was conducted at 50 °C for 24 hours with 5 RPM end-over-end mixing and TPA equivalents were estimated via fluorometric assay and compared to standards. All data reported represents the average of four technical replicates, and error bars represent standard deviation from the mean.

Assays on amorphous PET discs (**Figure 2B**) dictate a similar pH effect as the slurry, but with one intriguing difference; D2 CBM2a – Cutinase is clearly outperforming all constructs including the isolated CD at pH 6.0. Hole punched discs provide much less accessible surface area as compared to the PET slurry; thus, binding becomes more limiting. In most cases, CBM fusions produce a similar to slightly higher hydrolysis output as compared to the CD alone. At the pH optima, only the D2 CBM2a fusion is showing increased activity, with ∼2.1-fold higher TPA equivalent release. Better binding alone cannot explain this effect however, because this should be seen by other fusions as well. Instead, it is likely a cumulative impact where binding, and some other improvement not being directly tested here such as structural or thermal stability are providing unique benefits for this specific construct only. Interestingly, a similar trend was observed for this specific D2 CBM fused to a cellulase catalytic domain as well. On cellulose, the D2 CBM behaved like an outlier, at only it’s pH optima it showed the highest hydrolytic activity compared to the rest of the supercharged CBM library. However, binding appeared to play a more dominant role in that work, thus its effect on thermostability was not examined in detail.^31^ Additionally, this same D2 CBM2a – cellulase fusion was the best synergistic partner when paired with other enzymes for synergistic substrate deconstruction.^37^ Although binding seems to be a smaller player in the PET system, the distinct gains seen for the D2 CBM2a provides this specific construct with the unique opportunity to increase PET hydrolysis performance.

When correlating activity data to each construct’s net charge at each pH (**Figure 3**) for the CBM fusions only, no clear gaussian distribution indicative of a Sabatier-like trend is observed. Instead, higher activity is leftward shifted, favoring a negative net charge between -15 to -10 on both the PET slurry (**Figure 3C**) and amorphous disc (**Figure 3D**). On a per trial basis for milled PET (**Figure 3A**), the distribution of activity clearly trends to this negative charge. There is modest activity near a slightly positive charge of around 5, and this peaks around -10 before quickly falling off. On discs however (**Figure 3B**), D2 CBM2a is the clear outlier where its significant improvement is isolated from the general data trend. This behavior mimics the construct’s behavior on cellulose^31^ and further implies that there are other critical factors besides binding alone that can explain this improvement in activity.

**Figure 3.**
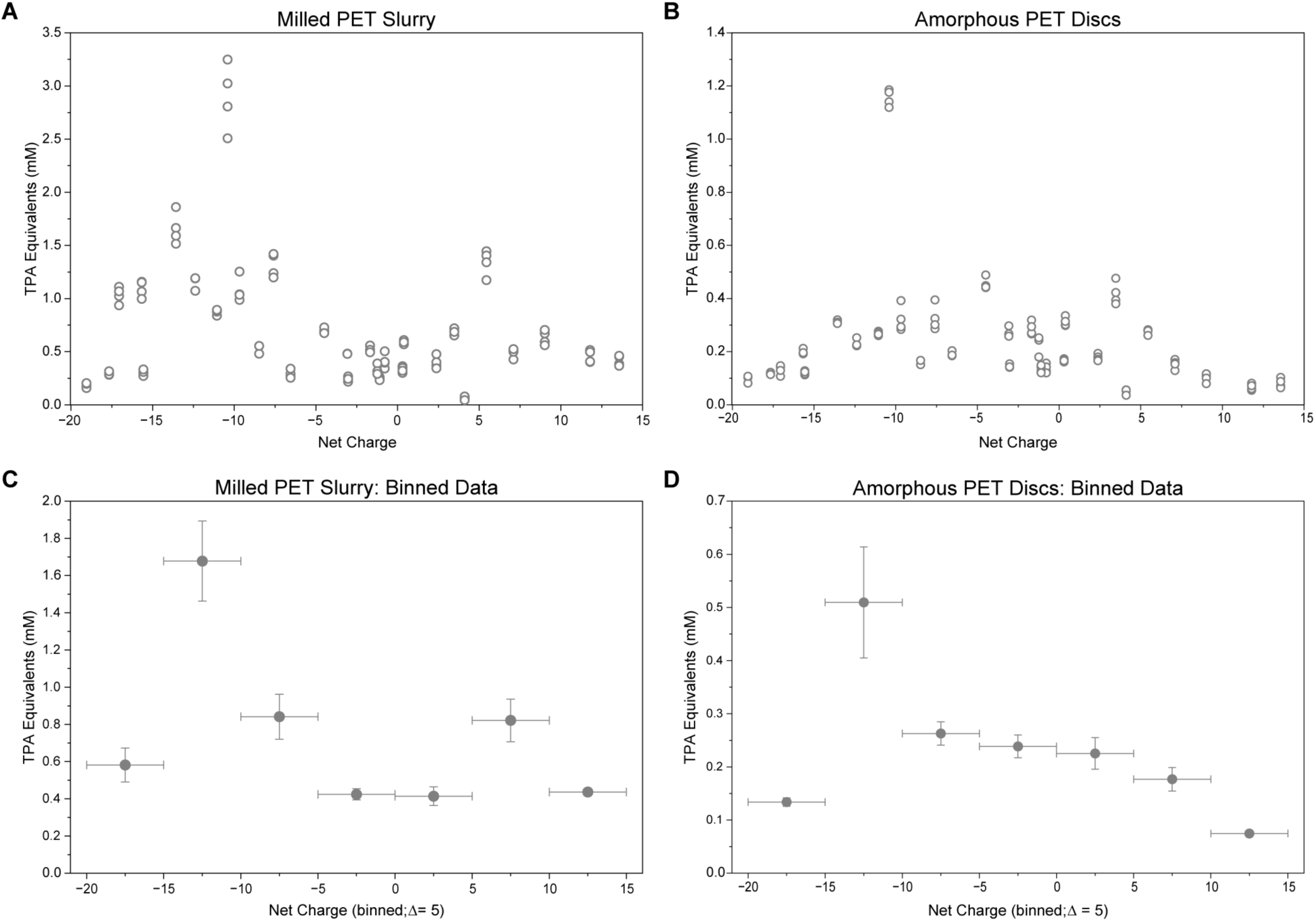
Chimeric CBM2a-cutinase enzyme activity does not show a strong Sabatier-like correlation with net charge. Relationship between CBM net charge and PET hydrolysis activity. (A) Individual trial hydrolysis measurements on milled PET originally shown in Figure 2A correlation with respect to net charge. (B) Individual trial hydrolysis measurements on amorphous PET discs originally shown in Figure 2B correlation with respect to net charge. (C) Binned hydrolysis activity data on milled PET slurry grouped by CBM net charge (binned ±5 charge units). (D) Binned hydrolysis activity data on amorphous PET discs grouped by CBM net charge (binned ±5 charge units). Hydrolysis activity is reported as TPA equivalents (mM). Y-error bars in binned datasets represent standard error of the mean for grouped measurements, and x-error bars denote bin width. No strong correlation between CBM net charge and PET hydrolysis activity was observed, despite marked differences in electrostatic surface properties between constructs.

### D2 CBM2a – Cutinase exhibits improved structural and functional thermostability

Hydrolysis assays across different buffer pH conditions suggested that electrostatic binding interactions likely play a more limited role in improving PET hydrolysis than initially hypothesized. Importantly, these assays identified D2 CBM2a – Cutinase as a clear standout for reasons beyond binding alone. To further investigate why this construct consistently outperformed the other fusions, thermostability of the entire library was assessed using differential scanning fluorimetry (DSF) across the same pH range used in the previous activity assays. Melt curves for the entire library are shown in **Figure 4** and illustrate global unfolding behavior and pH-dependent thermal transitions. **Figure 5** depicts the negative derivative of these curves, allowing direct quantification of melting temperatures. DSF was unable to resolve the melting temperature of the D1 CBM2a fusion due to aggregation-associated dye trapping. DSF data for this construct is provided in **Supplemental Figure S2**.

**Figure 4.**
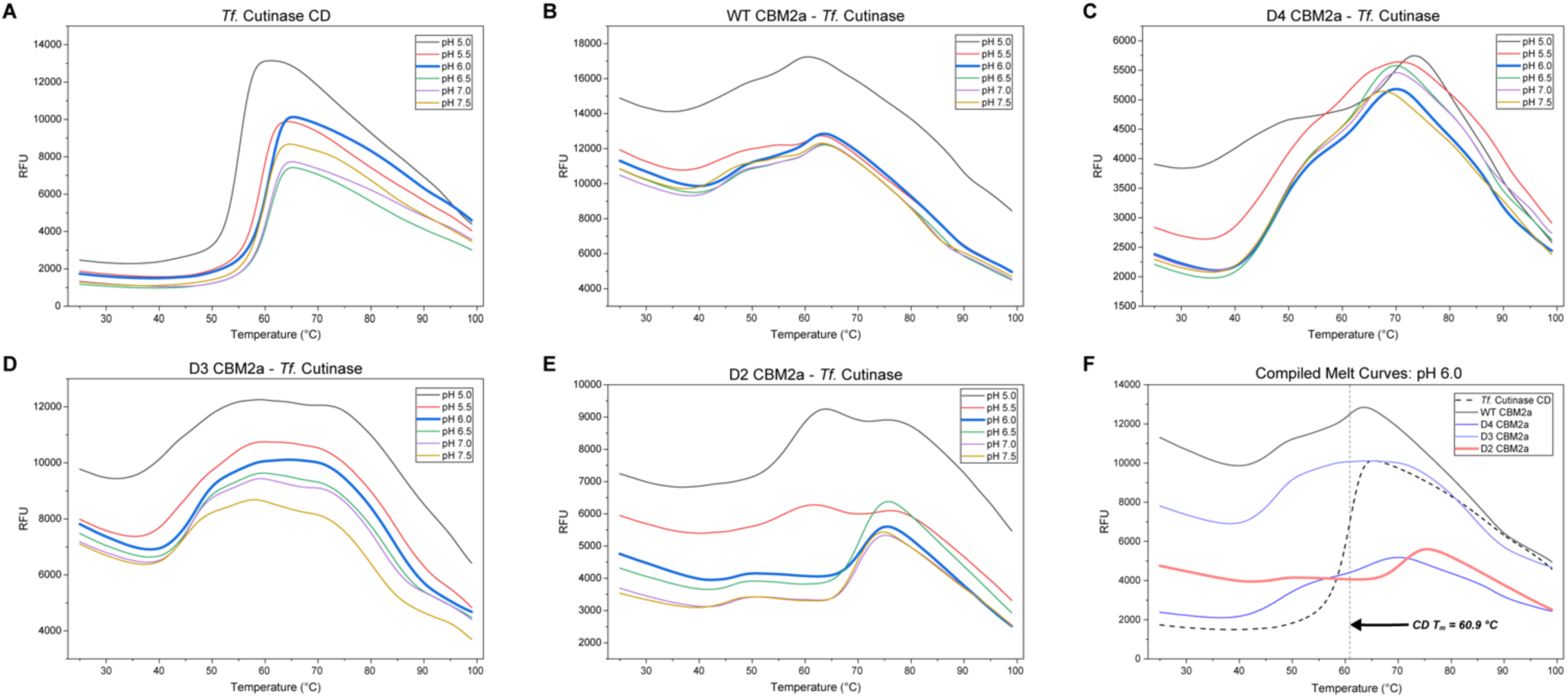
Differential scanning fluorimetry (DSF) melt curves of the Cutinase CD and charge-engineered CBM2a-Cutinase fusion constructs across varying buffer pH conditions. DSF melt curves for the isolated *T. fusca* Cutinase catalytic domain (A), WT CBM2a–Cutinase fusion (B), D4 CBM2a–Cutinase fusion (C), D3 CBM2a–Cutinase fusion (D), and D2 CBM2a–Cutinase fusion (E) collected across buffer pH conditions ranging from pH 5.0–7.5. Fluorescence intensity is plotted as a function of temperature. Curves illustrate global unfolding behavior and pH-dependent thermal transitions of each construct. The isolated catalytic domain exhibits a canonical single-domain melt transition, while CBM fusion constructs display more complex unfolding behavior associated with independent or cooperative melting of the CBM and catalytic domains. (F) Overlay of representative melt curves at pH 6.0 highlighting differences in unfolding behavior between constructs. The D2 CBM2a fusion exhibits a pronounced rightward shift in catalytic domain unfolding compared to the isolated catalytic domain, corresponding to an increase in apparent melting temperature. All proteins were tested in quadruplicates and data was processed in MATLAB to average replicates, subtract background fluorescence of a water blank, and smooth curves.

**Figure 5.**
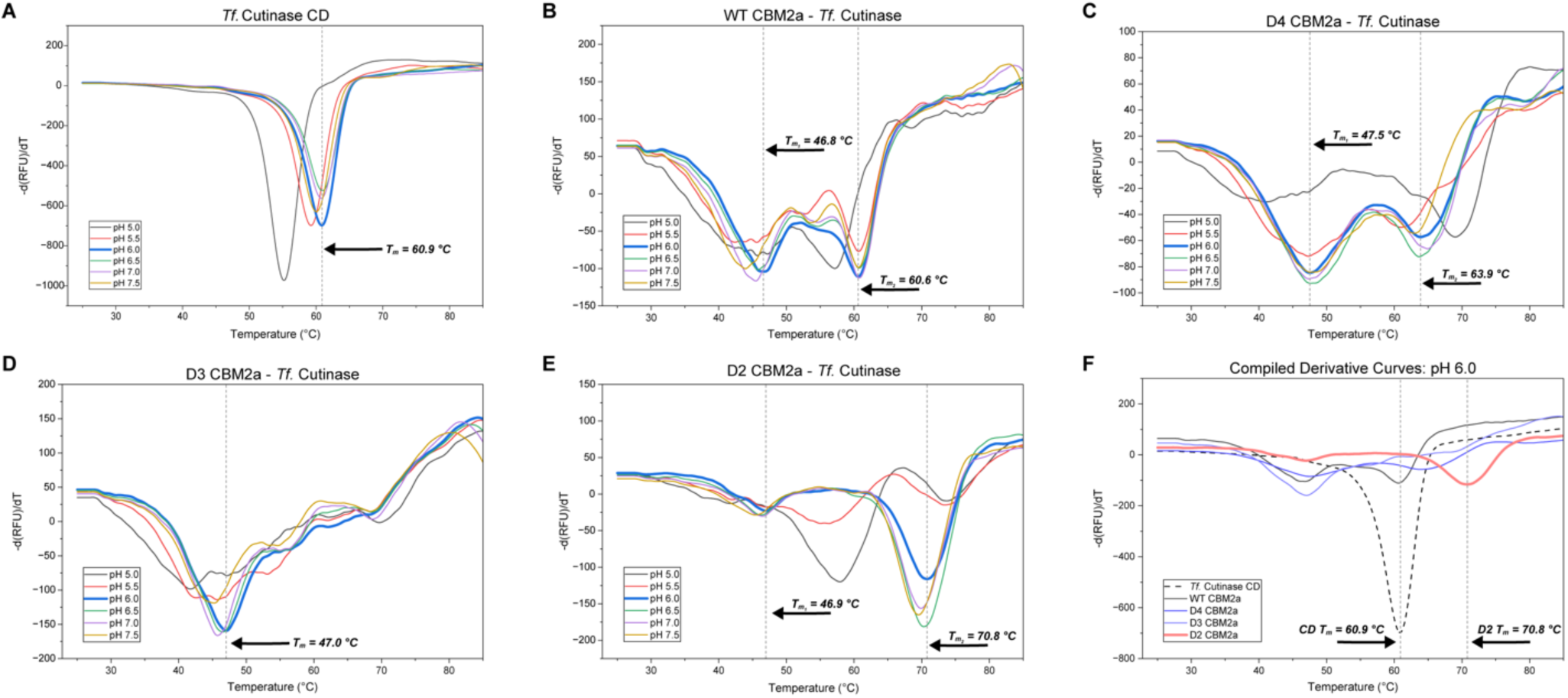
Derivative DSF curves used to quantify melting transitions of charge-engineered CBM2a-Cutinase fusion constructs. Derivative DSF curves for the isolated *T. fusca* Cutinase catalytic domain (A), WT CBM2a–Cutinase fusion (B), D4 CBM2a–Cutinase fusion (C), D3 CBM2a–Cutinase fusion (D), and D2 CBM2a–Cutinase fusion (E) collected across buffer pH conditions ranging from pH 5.0–7.0. Negative peaks correspond to thermal unfolding transitions and were used to determine apparent melting temperatures (*T_m_*). The isolated catalytic domain exhibits a single dominant melt transition, while CBM fusion constructs display multiple transitions associated with independent or cooperative unfolding of the CBM and catalytic domains. (F) Overlay of representative derivative melt curves at pH 6.0 highlighting shifts in catalytic domain unfolding between constructs. The D2 CBM2a fusion exhibits a pronounced rightward shift in catalytic domain melting transition relative to the isolated catalytic domain, corresponding to an approximately 10 °C increase in apparent melting temperature. Data was processed in MATLAB by first averaging the four replicates per melt curve and subtracting background fluorescence measured with a water blank. The negative derivative was calculated using the MATLAB gradient function and minima were identified with the min function in MATLAB.

The isolated Cutinase CD (**Figure 4A**) exhibits a canonical single-domain melt profile, with a clear pH dependence on melt behavior. Interestingly, the highest melting temperature (60.9 °C) is observed at pH 6.0 (**Figure 5A**), the same, pH where optimal catalysis was observed. This same trend is observed for all constructs suggesting that this enzyme may be more limited by stability than binding and is likely why a strong charge-activity correlation is not observed. For the native WT CBM2a fusion (**Figure 4B, 5B**), two melt transitions are observed, corresponding to a strong CBM melt event around 46.8 °C, and CD melt around the same temperature of the isolated CD. The most positive CBM fusion, D4 CBM2a (**Figure 4C, 5C**) exhibits similar melt behavior to the native fusion as well. For both enzymes, there is evidence of significant destabilization in DSF data at lower pH where several different melt events seem to be occurring, with noisy DSF traces. The D3 CBM fusion (**Figure 4D, 5D**) shows interesting melt behavior comparatively. Instead of two clear melt events, a broader hump is present in the melt trace, hinting that the initial lower temperature unfolding of the CBM is destabilizing the overall fusion, with both domains seemingly melting cooperatively. A similar destabilization was observed when this same CBM was fused to cellulase Cel5A^31^, but it destabilized the full-length enzyme to a much lesser degree than what was observed here. For the D2 CBM2a fusion (**Figure 4E, 5E**), DSF curves illustrate why this construct exhibited the best performance in PET hydrolysis. Despite showing strong destabilization at pH 5.0 and 5.5, a much smaller CBM melt event is observed at pH 6.0 onwards, as well as a rightward shift in the CD melt corresponding to a melt temperature of 70.8 °C, roughly 10 °C higher than the catalytic domain (**Figure 4F, 5F**). It seems therefore, that it is in fact improved thermostability, and not binding alone that produces the additive gain of function for this construct. A similar effect was observed for supercharged CBMs fused to an exo-active cellulase Cel6B where independent melting of a CBM seemed to decouple from the full-length construct melt, resulting in an increased catalytic unit melt and higher overall activity.^37^

Melt data does not suggest that the D2 CBM itself is thermostabilized. Additionally, when fused to a cellulase CD, this construct did not show any meaningful change in melt behavior, nor did circular dichroism suggest alteration of the CBM fold.^31^ There is more nuance to this effect, where this specific combination of CBM and Cutinase domain is behaving in such a manner that the overall fusion is stabilized. It is possible that the initial CBM melt may be providing enhanced flexibility, preventing CD unfolding, or that slight unfolding of the negatively charged domain may be binding to and stabilizing positive patches on the Cutinase surface to produce this effect. Further structural characterization will be required to determine the precise intramolecular interactions responsible for the apparent stabilization observed in the D2 fusion construct.

Although DSF provides insight into structural unfolding behavior, melting temperature alone does not elucidate catalytic competence following thermal challenge. Thus, residual activity assays were performed to assess whether the enhanced structural stability observed for D2 translated into improved functional persistence during PET hydrolysis (**Figure 6**). Enzyme aliquots were preincubated at increasing temperatures prior to hydrolysis assays, and remaining activity was quantified relative to untreated controls to determine T_50_ values. Results on milled PET (**Figure 6A**) depict progressive decreases in activity for the CD as pre-incubation temperature increases, followed by a significant drop in activity for the catalytic domain between 55 and 60 °C approaching the melting temperature measured with DSF. The D2 fusion retains substantial activity up to 55 °C, only decreasing thereafter, followed by a significant drop between 65 and 70 °C. This same trend holds consistently on amorphous discs (**Figure 6B**) for both enzymes, importantly indicating that, not only does D2 CBM2a exhibit a higher melting temperature, but this translates to a more robust catalytic performance under thermal stress. T_50_ values corresponding to the temperature where each enzyme loses half of its activity under thermal challenge corroborates this, identifying an 11.6 °C and 7.2 °C increase on PET slurry and discs respectively for the D2 CBM2a fusion. Tabulated melting temperatures for the entire library are displayed in **Table T1** and a summary of T_50_ values for D2 CBM2a compared to the Cutinase CD are depicted in **Table T2**. Together, these findings strongly suggest that enhanced catalytic persistence under thermal stress, rather than binding alone, is a dominant contributor to the improved PET hydrolysis performance observed for the D2 CBM2a fusion.

**Table T1.** Tabulated melting temperatures measured with DFS at pH 6.0. Melting temperature correspond to data shown in Figures 4 and 5. The D2 CBM2a fusion exhibits a near 10 °C increase in melting temperature compared to the Cutinase CD.

| Construct | CBM Charge | $T_m$ (°C, pH 6.0) | $\Delta T_m$ vs. CD (°C) |
| --- | --- | --- | --- |
| Cutinase CD | N/A | 60.9 | N/A |
| WT CBM2a | -2 | 60.6 | -0.3 |
| D4 CBM2a | 8 | 63.9 | +3.0 |
| D3 CBM2a | 6 | 47.0 | -13.9 |
| **D2 CBM2a** | **-8** | **70.8** | **+9.9** |

**Table T2.** Functional thermostability (T_50_) of the isolated *T. fusca* Cutinase catalytic domain and D2 CBM2a–Cutinase fusion on PET substrates. T_50_ temperatures corresponding to the temperature at which residual activity after thermal challenge is half that of unheated enzyme. Data presented in this table corresponds to logistic curve fits for the data presented in Figure 6. Curve fitting was done in Origin data analysis software. Curve fits can be viewed in Supplemental Figure S3.

| Construct | Substrate | $T_{50}$ (°C) |
| --- | --- | --- |
| Cutinase CD | Milled PET | 53.8 ± 1.12 |
|  | PET Discs | 56.6 ± 1.37 |
| D2 CBM2a | Milled PET | 65.4 ± 0.62 |
|  | PET Discs | 63.8 ± 1.01 |

**Figure 6.**
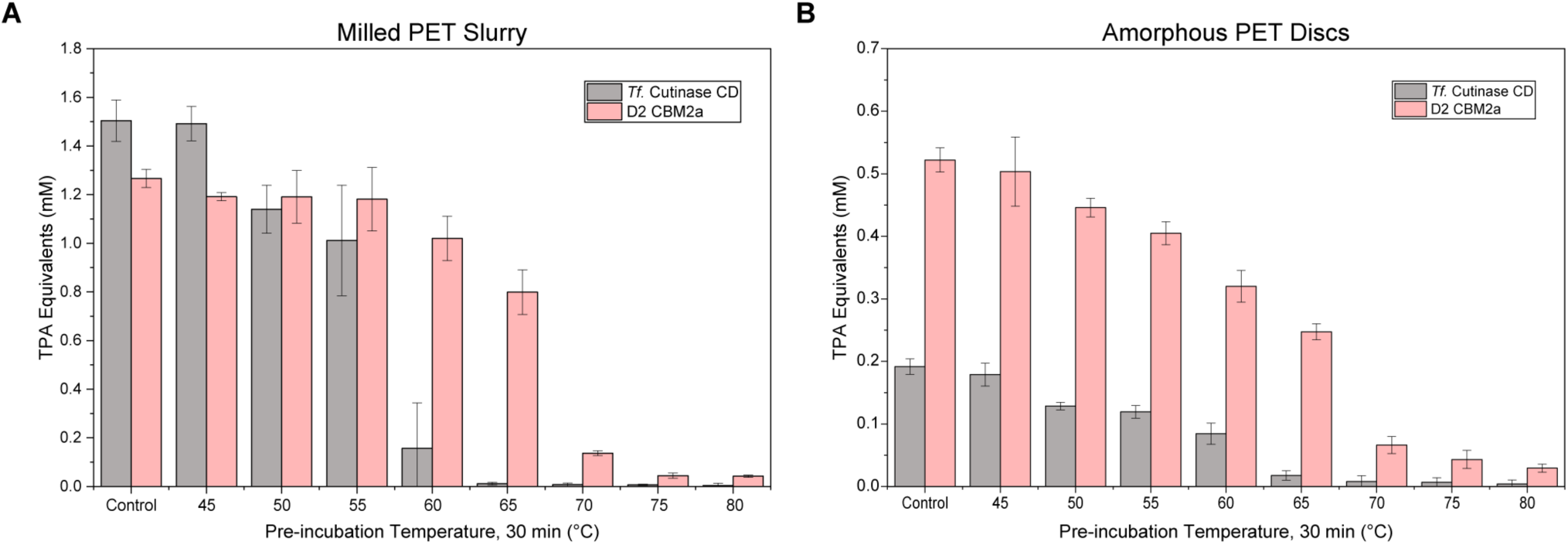
Functional thermostability (T_50_) analysis of the D2 CBM2a – Cutinase fusion compared to the isolated Cutinase CD shows improved functional persistence after preincubation thermal challenge. The isolated *T. fusca* Cutinase catalytic domain (CD) and D2 CBM2a–Cutinase fusion were pre-incubated for 30 min at the temperatures indicated on the x-axis prior to PET hydrolysis assays. Residual hydrolysis activity is reported as TPA equivalents (mM) measured after incubation with milled PET slurry (A) or amorphous PET discs (B). Enzymes diluted in buffer were aliquoted into PCR tubes (60 µL) before thermal challenge. Residual activity was tested by adding 50 µL of clarified soluble supernatant after heat treatment to 4 mg of substrate (120 nmol enzyme/g substrate), with 50 mM sodium phosphate, pH 6.0 and incubated for 24 hours with 5 RPM end-over-end mixing at 50 °C. The D2 CBM2a fusion retained substantially greater hydrolytic activity following thermal challenge compared to the isolated catalytic domain, particularly at elevated temperatures. These data demonstrate that the D2 fusion exhibits improved functional persistence and thermal tolerance on both PET substrates. All data represents the average of four trials, and error bars represent standard deviation from the mean.

### Surface charge modulates PET binding behavior

Fusion of CBM binding partners did not produce the expected outcome in overall PET hydrolysis for the entire library; improvements appear to be driven primarily by thermostabilization of the key D2 construct. Supercharging was a potent method to modulate binding to cellulose for these same CBMs^31^, but polyesterase activity does not adequately clarify if this same phenomenon exists on PET. To probe whether or not the supercharging design principle does produce marked differences in PET binding, CBMs were fused to green fluorescent protein (GFP) and used to assess PET binding through pull-down binding assays. It is important to note that this scheme has substantial limitations on PET. GFP tagged CBMs showed low overall binding to PET, and at higher concentrations, the signal-to-noise ratio became too large to reliably distinguish meaningful binding trends. Additionally, there was strong evidence of nonspecific binding, where GFP alone exhibited substantial PET association under all conditions. Additionally, the presence of BSA significantly altered apparent protein partitioning behavior, particularly on hydrophobic PET discs, suggesting that interfacial and wetting effects strongly influence measured depletion signals.^42–44^ For these reasons, full isotherms were not constructed, and binding only characterized based on GFP depletion at one fixed concentration.

**Figure 7** depicts these binding assays on milled PET (**A, C**) and PET discs (**B, D**) both with 0.05% BSA (**C, D**) and without BSA present (**A, B**). The high depletion observed for untagged GFP strongly suggests a high propensity for protein to adsorb to the PET surface, and this tracks with previous literature identifying that PET hydrolases efficiently adsorb to the PET surface in a monolayer, primarily driven by nonspecific interactions.^45^ For assays without BSA present, it is not possible to rule out potential background adsorption to the microplate itself, however, noticeable differences are observed for the different CBMs. On milled PET (**Figure 7A**), the most positive CBM (D4) shows the highest GFP depletion for fusion constructs, with slight decreases tracking as charge becomes more negative with D2 CBM2a. WT CBM2a however shows the lowest overall PET binding, potentially arising from different background effects. After this initial binding step, samples were re-equilibrated with fresh buffer to probe redistribution behavior after dilution. Recovery values (right y-axis) near zero indicate little redistribution beyond dilution, positive values indicate recovery of the fluorophore in solution indicative of desorption from PET, and negative values indicate fluorescent loss related to additional PET association after dilution. For the CBM fusions, only D2 CBM2a shows slight net fluorescence recovery after dilution, while both positively supercharged CBMs (D4 and D3) continue exhibiting net PET association after re-equilibration. Interestingly, GFP alone exhibited both the largest initial depletion and fluorescence recovery. This result seems to indicate that nonspecific adsorption between GFP and PET is not only strong, but also dynamic rather than fully irreversible.

**Figure 7.**
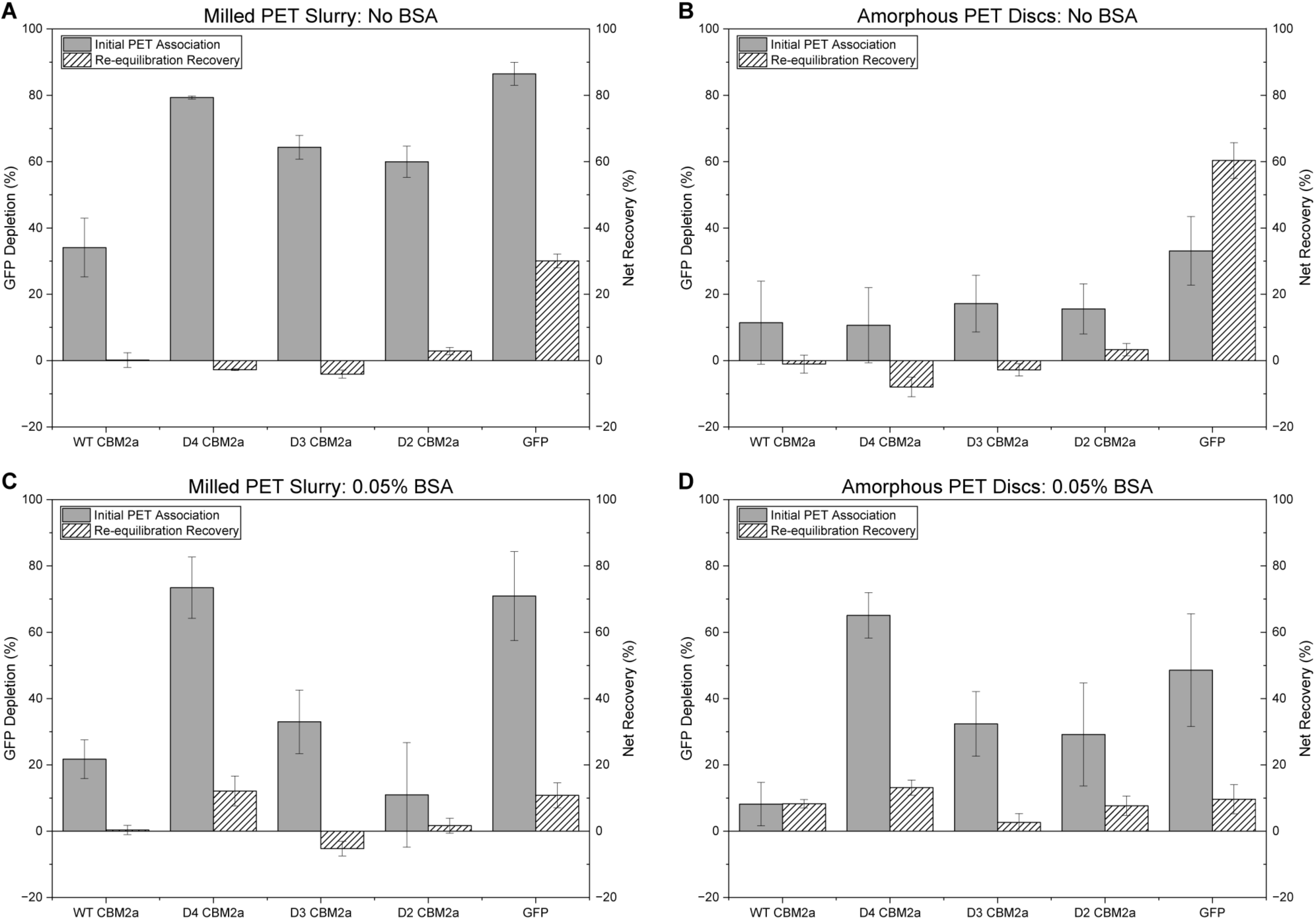
Pull-down GFP tagged CBM binding assays shows a stronger PET adsorption for positively charged CBMs, but strong non-specific adsorption for untagged GFP. Binding interactions between GFP-tagged CBM2a variants and PET substrates were assessed through pull-down depletion assays using milled PET slurry and amorphous PET discs. GFP depletion (%) from solution was used as a proxy for substrate association. Assays were performed on milled PET slurry without BSA (A), amorphous PET discs without BSA (B), milled PET slurry with 0.05% BSA (C), and amorphous PET discs with 0.05% BSA (D). Initial binding measurements are shown alongside re-equilibration binding measurements following replacement with fresh buffer. A total 4 mg of PET was used in binding wells with protein concentration fixed at 1 µM. 50 mM HEPES buffer pH 6.5 was used for binding. GFP depletion was recorded by comparing the difference in fluorescence between binding wells and standard wells containing no substrate. GFP alone exhibited substantial nonspecific PET association, particularly on milled PET slurry. The highly positively charged D4 CBM2a variant showed the greatest PET depletion overall, while the D2 CBM2a fusion exhibited comparatively weaker PET association despite displaying superior hydrolysis performance in catalytic assays. These results suggest that increased PET adsorption alone does not directly translate to improved PET hydrolysis efficiency. Data points represent the average of six replicates and error bars represent standard deviation from the mean.

On amorphous discs (**Figure 7B**), a generally low overall initial binding is recorded, confirming there is a much lower accessible surface for CBMs to bind to on this substrate. This effect is likely the root cause producing the low overall activity observed for most constructs on PET discs. Interestingly, the D4 CBM shows a larger bound portion on PET, suggesting that the increased positive charge likely does improve substrate adhesion, but does not appear to translate efficiently into productive hydrolysis under dilute conditions. Re-equilibration data for the CBM fusions depict a strong correlation between recovery and net charge where the most positively charged constructs exhibit continued PET association after dilution, while the more negatively charged constructs exhibit fluorescence recovery associated with desorption into solution. Although to a much lower degree than on milled PET, GFP still exhibits significant nonspecific adsorption to the PET discs, with an even larger fraction redistributing into solution upon dilution.

The addition of BSA partially suppresses nonspecific background interactions with assay interfaces and more clearly illustrates electrostatic trends. On milled PET, there is a clean correlation where the most positive CBM exhibits the highest overall PET binding (**Figure 7C**), while the most negative construct shown here, D2 CBM2a exhibits the lowest. For GFP, a substantial portion both binds to PET and redistributes into solution after dilution, confirming that nonspecific adsorption to PET does indeed play a substantial role in this system. Key trends related to PET discs generally stay the same with BSA present (**Figure 7D**), with the clear point being positively charged CBMs do bind PET more effectively than native, or negative counterparts. Increasing positive charge appears to enhance PET association, hence the increased effect observed comparing the most positive domain D4 to D3 which is positively supercharged to a lesser degree. These results indicate that the reasoning in formulating the initial hypothesis does indeed hold true. It was correctly hypothesized that positive charge would increase PET adsorption, however, this trend did not correlate with hydrolysis output. These findings highlight the mechanistic nuance within this system; increasing bound enzyme is not sufficient to increase hydrolytic activity as initially thought, and observed on cellulose.^31^ While supercharging does improve PET association, productive PET hydrolysis is not governed by maximal surface adsorption. While D4 CBM2a has the highest propensity to bind PET, it significantly underperforms both the CD and D2 CBM2a fusion. In fact, D2 CBM2a only weakly associates with PET at several conditions, despite being the most active fusion on PET. These assays further indicate that PET is highly prone to dynamic, nonspecific protein adsorption, and this is likely why the CD can still effectively degrade PET without a binding unit. Based on these findings, although supercharging can improve PET association, this is not the strongest route for increasing hydrolytic efficiency of PET hydrolases. Instead, thermostabilization, as observed with D2 CBM2a, appears to be a more effective route for improving PET hydrolysis, despite emerging as an unintended outcome of this study.

### Thermostabilized D2 CBM2a – Cutinase fusion exhibits enhanced PET hydrolysis at elevated temperatures

Despite the relatively modest differences in PET hydrolysis arising from appended binding modules alone, the thermostabilization observed for D2 CBM2a–Cutinase provides a potential avenue to exploit higher hydrolytic efficiency through elevated reaction temperatures. Increased temperatures provide two key advantages: i) catalytic reaction rates increase despite accelerated enzyme deactivation, and ii) PET becomes more amenable to enzymatic hydrolysis. On the latter point, PET exhibits a crystallinity-dependent glass transition temperature (*T_g_*) ranging from 65–80 °C, with highly crystalline PET approaching the upper end of this range.^46^ As hydrolysis temperatures approach T_g_, water acts as a potent plasticizer, increasing polymer chain mobility and facilitating deeper penetration into the bulk substrate, thereby improving accessibility for enzymatic hydrolysis.^10,11,47,48^ For these reasons, the thermostabilization observed for D2 CBM2a–Cutinase is particularly advantageous for PET hydrolysis under elevated reaction temperatures.

To assess whether the observed structural thermostabilization translates to improved hydrolysis performance at elevated temperatures, the D2 CBM2a fusion and isolated Cutinase CD were incubated at several different temperatures, starting from the CD optimum (50 °C) for 4 hrs, and 24 hrs reaction time (**Figure 8**). Interestingly, after four hours of incubation with milled PET (**Figure 8A**), the catalytic domain exhibited a slightly higher hydrolysis yield as compared to the 50 °C optimum, but functional persistence becomes limiting, as accumulated enzyme loss results in a lower overall yield after 24 hrs (**Figure 8B**). For the D2 CBM2a fusion however, the improvements are immediately observed between 55 – 60 °C, exhibiting a 10 °C increase in optimal hydrolysis temperature. Improved catalytic persistence at 60 °C enabled a ∼3.4-fold increase in hydrolysis yield compared to the CD alone at its temperature optimum. Although the CD outperformed the D2 CBM2a fusion in initial screens at 50 °C, thermostabilization of the bulk ensemble construct observed for D2 CBM2a results in functional persistence at elevated temperatures where PET is more amenable to enzymatic degradation, thus, producing higher overall activity.

**Figure 8.**
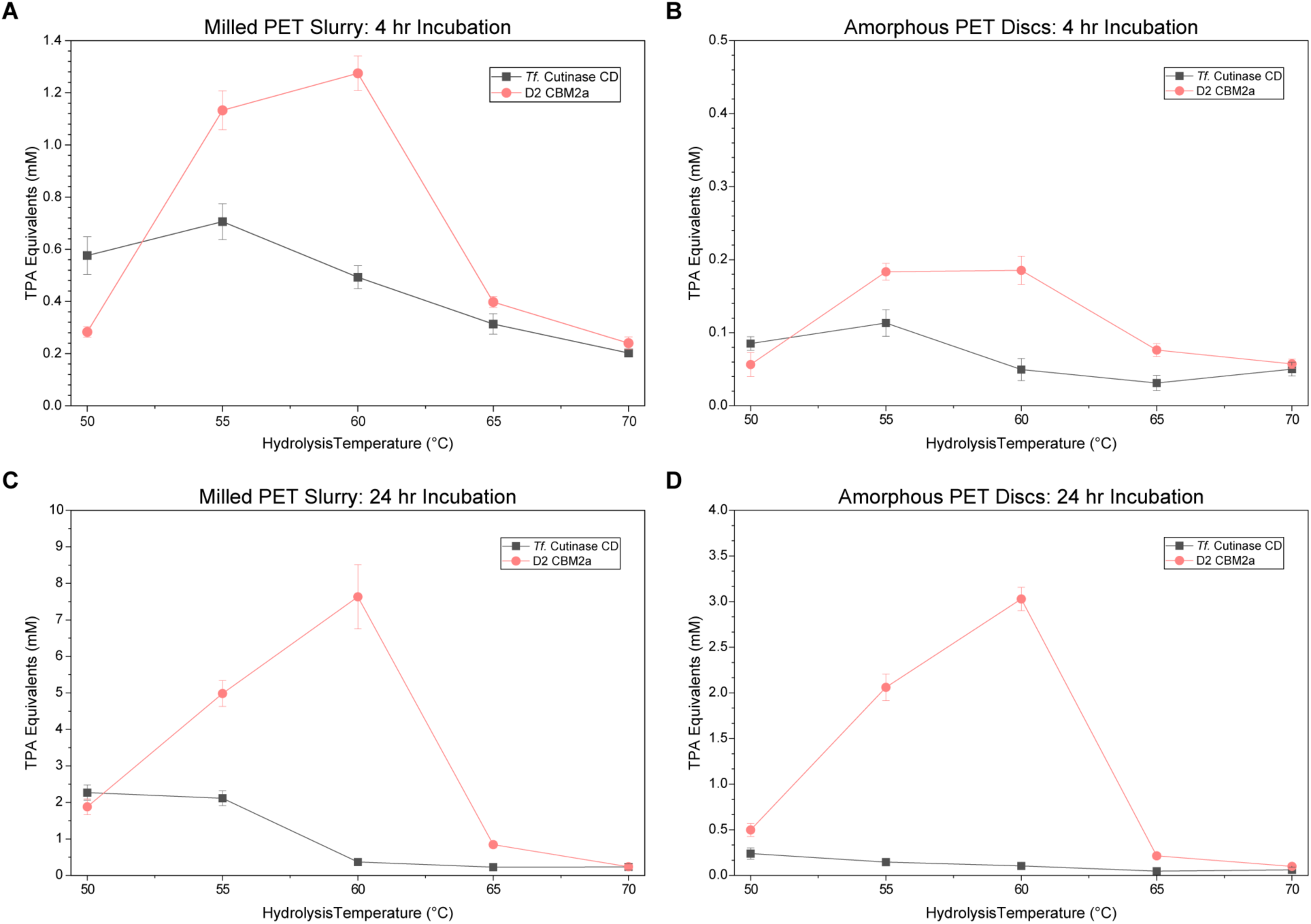
Sweeping hydrolysis temperature shows a 10 °C increase in D2 CBM2a – Cutinase optimal hydrolysis temperature. Hydrolysis activity of the isolated *T. fusca* Cutinase catalytic domain (CD) and D2 CBM2a–Cutinase fusion was assessed across a range of reaction temperatures using milled PET slurry and amorphous PET discs. Hydrolysis yields are reported as TPA equivalents (mM) following 4 h incubation with milled PET slurry (A), 4 h incubation with amorphous PET discs (B), 24 h incubation with milled PET slurry (C), and 24 h incubation with amorphous PET discs (D). Enzyme dilutions were prepared in water to supply a constant 120 nmol enzyme/ g substrate loading with 4 mg substrate in 50 mM sodium phosphate pH 6.0. Reaction plates were incubated at the temperatures indicated on the x-axis for their respective timepoints with 5 RPM end-over-end mixing. The isolated catalytic domain exhibited maximal activity near 50–55 °C followed by rapid loss of activity at elevated temperatures, consistent with thermal inactivation. In contrast, the D2 CBM2a fusion displayed a shifted optimal hydrolysis temperature near 60 °C and retained substantially greater hydrolytic activity under elevated temperature conditions. These results demonstrate that thermostabilization of the D2 fusion enables enhanced PET depolymerization at temperatures more favorable for PET chain mobility and hydrolysis. Data represents the average of four replicates, and error bars represent standard deviation from the mean.

These same general trends hold on amorphous discs at both four (**Figure 8B**), and 24 hr (**Figure 8D**) timepoints. Interestingly, the isolated CD shows a higher initial activity after four hours at 50 °C, but the overall hydrolysis yield is higher for D2 CBM2a after 24 hours, mimicking results seen in the initial pH sweeps. This result strongly suggests stability limitations in this native CD. On a qualitative measure, each construct, with the exception of D2 CBM2a exhibited some degree of insoluble pelleting after incubation with PET discs, likely attributed to protein aggregation and denaturation after long incubation. When incubated with a substrate with less surface accessibility, a greater portion of loaded enzyme will be present in solution, not engaged with substrate, where it may be prone to denaturation or aggregation. CBM fused cellulases have been shown to experience this problem, where substrate engagement and formation of a stable enzyme-substrate complex prevents denaturation.^31^ This effect is likely a major limiting factor in this work, hence the lower long-term activity of the CD. Stabilization of the D2 CBM2a fusion overcomes this limitation, resulting in a higher activity for D2 CBM2a at 50 °C, compounded with any beneficial binding gained with the fused CBM. This effect is amplified dramatically at the higher temperature optimum of D2 CBM2a (60 °C), where a ∼12.5-fold difference is observed compared to the isolated CD at 50 °C. It is important to note that while this an exciting metric for improvement, overall conversion is still rather low at roughly 2% conversion, and the fractional increase in activity is amplified by the low baseline activity observed by the CD.

To explore this effect further and exploit the functional gains in the D2 CBM2a fusion, both the thermostabilized fusion and Cutinase CD were incubated with PET for 72 hours at both 50 and 60 °C (**Figure 9**). At 60 °C, on both milled PET (**Figure 9A**) and amorphous PET discs (**Figure 9B**), the D2 CBM2a fusion shows the highest overall activity, whilst the isolated CD is inactivated, likely thermally denatured. At the end of this time-course, a final ∼1.83-fold increase in hydrolysis yield on milled PET, and ∼9.63-fold increase on PET discs is observed comparing the D2 fusion at 60 °C to the Cutinase CD at 50 °C where both enzymes have levelled off in activity. In this timeframe, the maximum conversion measured for D2 CBM2a was 10.05 ± 1.83% on milled PET, and 3.15 ± 0.466 % on amorphous PET discs, corresponding to a ∼4.6% and ∼3.0% increase compared to the isolated CD respectively. Importantly, the substantially larger enhancement observed for intact PET discs compared to milled PET may hold particular relevance for industrial PET depolymerization processes. Highly milled PET powders are experimentally useful as model PET substrates; however, particle size reduction introduces preprocessing and energy costs that may limit industrial practicality. For this reason, the activity improvement observed on intact PET substrates becomes increasingly important from an industrial perspective. The strong relative performance of the D2 CBM2a fusion on intact PET discs suggests that thermostabilized fusion constructs may provide greater advantages for the depolymerization of minimally processed PET feedstocks.

**Figure 9.**
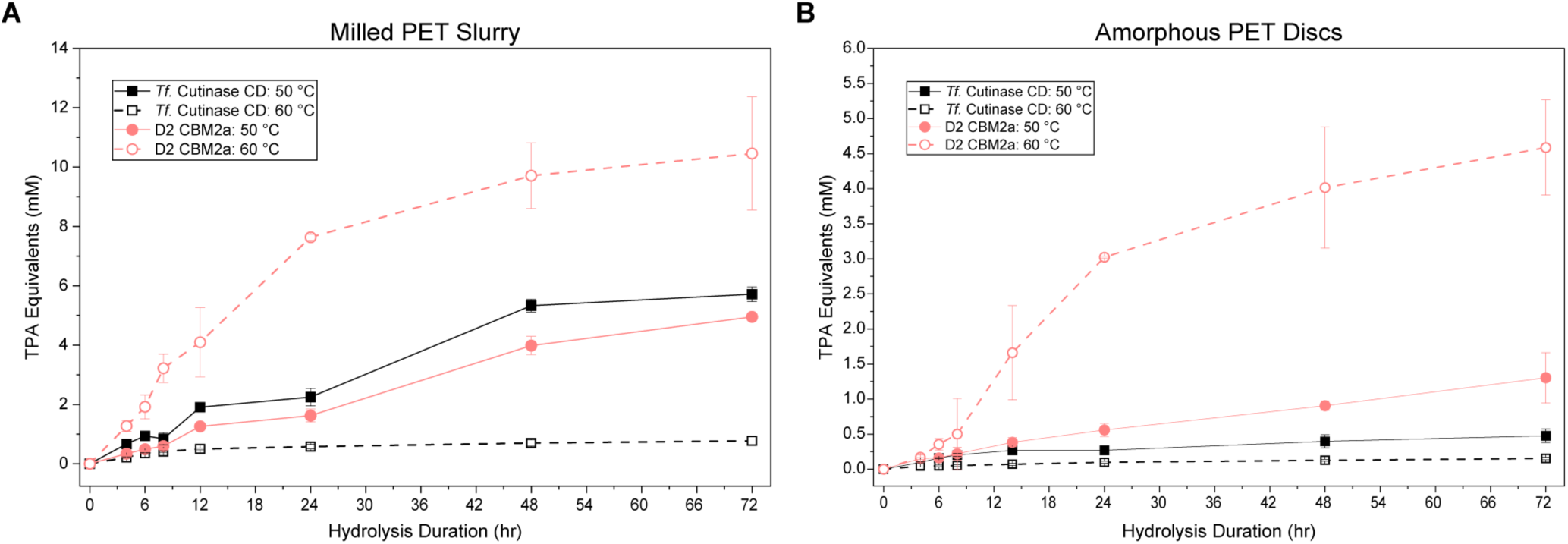
Extended time course PET hydrolysis performance for D2 CBM2a-Cutinase fusion compared to native *T. fusca* Cutinase catalytic domain at elevated temperatures. Time-dependent PET hydrolysis assays were performed using the isolated *T. fusca* Cutinase catalytic domain (CD) and D2 CBM2a–Cutinase fusion at 50 and 60 °C across a 0-72 h incubation period. Hydrolysis yields are reported as TPA equivalents (mM) for milled PET slurry (A) and amorphous PET discs (B). Assays were performed by incubated enzyme dilutions in water with 4 mg PET substrate at a constant loading of 120 nmol enzyme/g substrate in 50 mM sodium phosphate pH 6.0. The isolated catalytic domain exhibited limited activity at 60 °C, consistent with rapid thermal inactivation, while the D2 CBM2a fusion retained substantial hydrolytic activity throughout the incubation period at elevated temperature. The thermostabilized D2 fusion achieved the highest overall PET conversion on both substrates, demonstrating improved catalytic persistence under conditions favorable for PET chain mobility and depolymerization. Data points represent the average of four replicates and error bars denote standard deviation from the mean.

Although the D2 CBM2a fusion presents a stark improvement compared to the native enzyme, hydrolysis yield is capped near 10% conversion, limited by the poor baseline PET degrading activity of the *T. fusca* Cutinase enzyme. Similar trends have been observed in highly active engineered PET depolymerases where thermostabilization has enabled dramatic improvements in PET conversion. For example, Leaf-branch compost Cutinase (LCC) thermostabilized variant ICCG achieved near complete depolymerization (90%) within 10 hr following thermostabilization and active-site engineering.^16^ Native LCC already exhibits a substantially higher baseline PET hydrolysis activity (∼50% conversion) as well as higher thermostability (T_m_ = 84.7 °C), enabling the improved variant to reach near complete substrate depolymerization.^16^ The Cutinase enzyme used in this study exhibits comparatively poor intrinsic PET hydrolysis activity and thermostability, limiting absolute conversion despite similar fold-improvements in hydrolysis performance following thermostabilization.

### Substrate accessibility influences the hydrolysis advantage of D2 CBM2a–Cutinase

The dominant mechanism contributing to improved hydrolysis for D2 CBM2a appears to be enhanced thermostability, but to this point, the impact of CBM binding for the D2 construct is unclear. Data here has shown different behavior in situations where enzymes are presented a substrate with a large accessible surface (milled PET) compared to a more surface-limited configuration (hole punched PET discs). This effect was probed further in order to assess if there are marked differences in enzyme behavior as substrate loading changes (**Figure 10**) for both the isolated CD, and active D2 CBM2a fusion. These effects were only assessed at 50 °C as the Cutinase CD does not show activity at 60 °C, preventing meaningful comparison. On milled PET (**Figure 10A**), both enzymes seem to scale evenly with substrate loading. At 50 °C, the CD is more active than the fusion as seen previously, and for both enzymes, hydrolysis yield increases as more substrate is added. This result reinforces the hypothesis that the Cutinase CD is not limited in adsorbing to PET derived from prior pH sweeps and corroborated with GFP binding assays. On an accessible finely milled substrate with high surface area, there is little CBM advantage, and instead, activity scales near linearly with an increased amount of substrate present.

**Figure 10.**
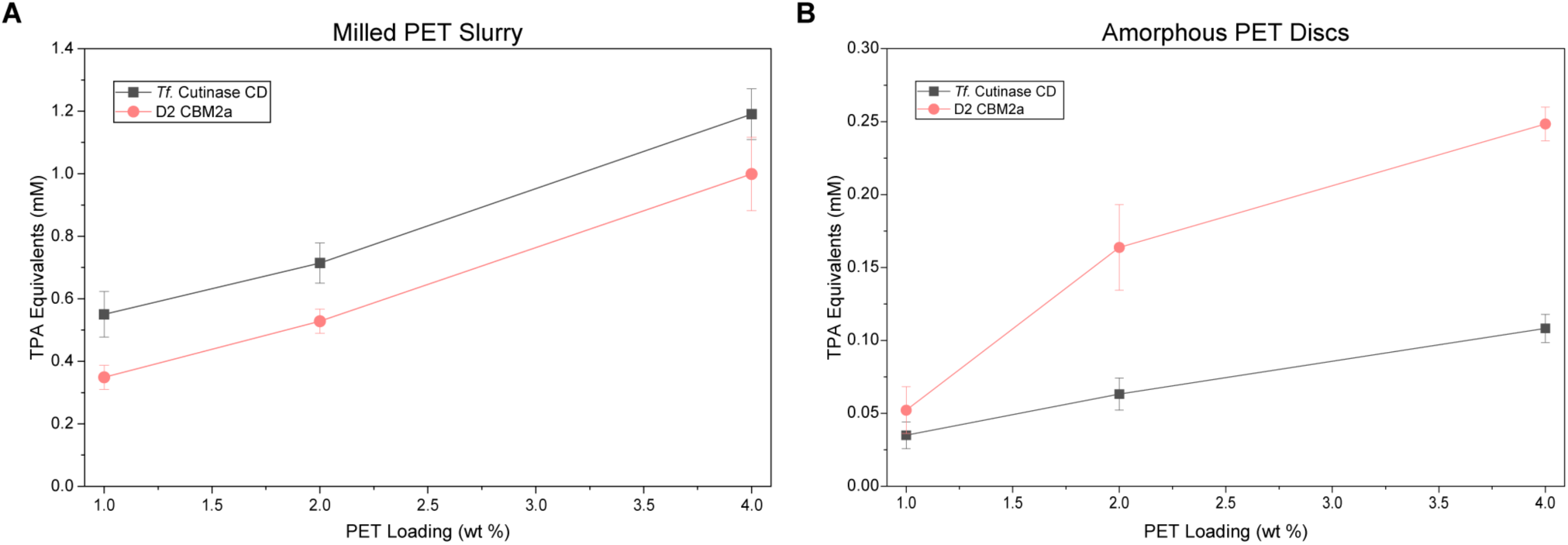
Influence of substrate loading on PET hydrolysis activity for D2 CBM2a – Cutinase fusion and isolated Cutinase CD. Hydrolysis activity of the isolated *T. fusca* Cutinase catalytic domain (CD) and D2 CBM2a–Cutinase fusion was assessed across increasing PET substrate loadings at 50 °C using milled PET slurry (A) and amorphous PET discs (B). Hydrolysis yields are reported as TPA equivalents (mM) following incubation with PET loadings ranging from 1–4 wt %. Enzyme loading was maintained at a constant 120 nmol/g of enzyme to explore effects of substrate loading alone. Assays were performed in 50 mM sodium phosphate, pH 6.0 and plates were incubated for 24 h with 5 RPM end-over-end mixing. On milled PET slurry, both constructs exhibited approximately linear increases in hydrolysis yield with increasing substrate concentration, with the isolated catalytic domain maintaining slightly higher overall activity. In contrast, the D2 CBM2a fusion displayed a disproportionately larger increase in hydrolysis activity on amorphous PET discs at low substrate loading, suggesting that CBM-mediated substrate interactions become more important under surface-limited conditions. These results indicate that the contribution of the appended CBM to PET hydrolysis is substrate-context dependent and secondary to the dominant thermostability-driven improvements observed for the D2 fusion construct. All data points represent the average of four replicates and error bars represent the standard deviation from the mean.

There is a marked difference on the more surface-limited PET discs for the D2 CBM fusion (**Figure 10B**). At each loading, a consistent higher activity is observed for the D2 CBM2a chimera, but more interestingly, the improvement scales with substrate load. This fusion shows a sharp increase from 1% to 2% PET solids loading, followed by a smaller gain from 2 to 4% while the CD shows a similar linear increase as the milled PET result tracking with increased hydrolysis yield due to a higher substrate concentration. This suggests that the CBM provides the greatest benefit under surface-limited conditions, but activity becomes constrained by another limiting factor at higher loading. Soluble pNP-Acetate hydrolysis assays did not show statistically meaningful differences in catalytic activity between the isolated CD and D2 CBM2a fusion, suggesting that the improved hydrolysis observed on PET discs is not driven by altered intrinsic catalytic turnover (**Supplemental Figure S4**). Thus, the CBM appears to contribute to PET hydrolysis in a substrate context dependent manner, but this contribution is subtle and secondary to the larger thermostability driven gains observed for D2. Similar conclusions have been drawn in other works that have attempted to fuse native CBMs to an LCC variant, where CBM gain was only observed at loadings below 5% loading, thereafter the binding contribution to improved activity is nullified.^28^ As mentioned earlier, this LCC catalytic domain is much more stable and exhibits a higher baseline PET activity, so it is highly probable that the CBM binding contributions at low loading are more visible when paired to this more active and thermostable CD. In this system, stability still seems to be the key limiting factor, and although the appended CBM does seem to provide some beneficial contribution, stabilization of the *T. fusca* Cutinase CD is much more effective in improving PET hydrolysis.

## CONCLUSION

In this study we have successfully identified a novel charge engineered CBM – PET hydrolase chimera that exhibited superior thermostability and substantially improved activity on various PET substrates compared to the native *T. fusca* Cutinase catalytic domain (CD). The family-2a CBM natively appended to a Cel5A cellulase found in *T. fusca* was engineered in our previous work to generate several constructs spanning a broad net charge range that previously exhibited superior cellulose binding characteristics and resulting activity as a result of the charge engineering.^31^ Drawing similarities between PET and cellulose on the basis of substrate electrostatics and planar aromatic backbones in the molecular structure, we had hypothesized that positively charged CBMs would show superior PET binding, thus potentially increasing PET hydrolysis rate through improved substrate targeting and recognition, if this enzyme-substrate system was adsorption limited. While positive CBMs did show stronger binding to PET in GFP based pull-down binding assays, this did not produce a resulting improvement in activity. Instead, a slightly negative charged CBM fusion, D2 CBM2a – Cutinase, was the only fusion construct to display any competitive improvement in activity, despite exhibiting poorer PET binding than its positive charge counterparts. This construct exhibited activity comparable to the isolated Cutinase CD on milled PET, as well as a ∼2-fold improvement in hydrolysis yields on amorphous PET discs, hinting that binding may not be the most important limiting factor in this system.

Probes into the cause of this change identified gains in structural stability for the D2 CBM2a fusion only, with a 10 °C increase in melting temperature of the ensemble chimera at pH 6.0 based on DSF. This increase in the D2 CBM fusion’s melt temperature translated to improved functional persistence post thermal challenge, an increase in optimal hydrolysis temperature, and higher time dependent yields at this higher temperature. At these higher temperatures, this fusion was able to achieve roughly 10% conversion of PET to TPA, a 4.6% absolute increase compared to the isolated catalytic unit. The exact driving force for this thermostabilization is not fully understood, however. The D2 CBM itself does not exhibit some inherently higher melt transition, showing strong pH dependent melt events comparable to the other CBMs examined here. Previous work with this CBM did not show a similar effect when fused to a completely different enzyme catalytic domain,^31^ thus unique intramolecular interaction between this specific CBM and the Cutinase CD must be driving this result. As a result of charge engineering, it is likely that the initial melt of the D2 CBM may produce some flexible region that may interact with charge complementary regions on the CD, stabilizing the overall structure and altering temperature dependent unfolding pathways. Supercharged CBMs have been previously shown to significantly alter thermal unfolding pathways. Previous work identified a similar effect for a fusion enzyme that natively exhibited cooperative melt initiated by the CBM. For one key supercharged CBM fusion, these melt transitions decoupled, with the catalytic unit exhibiting a higher melting transition than the native enzyme driving activity improvements.^37^ Although thermostabilization was not the intended goal in this work, it provided a potent mechanism for enzyme improvement for this PET hydrolase. These findings collectively suggest that enhanced catalytic persistence, rather than improved PET adsorption of enzymes alone, is the dominant factor governing the improved hydrolysis performance of the D2 fusion construct.

Although thermostabilization emerged as the dominant mechanism governing hydrolysis performance in this system, the appended CBM still appears to provide subtle, substrate-dependent contributions to PET depolymerization. Binding assays and substrate loading studies collectively suggest that CBM-mediated PET interactions are more important under surface-limited conditions, where productive engagement may become a limiting factor for the Cutinase catalytic domain. Binding assays indicated strong a propensity for non-productive engagement with a highly surface accessible substrate like milled PET, and as such, hydrolysis assays are overwhelmingly skewed towards the isolated catalytic domain at normal operating temperature. Thus, binding did not appear to be a limiting factor on this substrate. With a more surface-limited substrate like PET discs, CBM contributions are more apparent in the D2 CBM fusion, with a nonlinear increase in activity with increasing substrate load at low overall loading. However, the most positive D4 CBM that exhibited stronger PET binding did not comparatively show greater activity. The failure of this most positive construct to enhance PET discs hydrolysis suggests that increased adsorption alone is insufficient. Instead, it may promote nonproductive surface association where the CBM binds PET in non-catalytically conformations. Thus, D2 likely succeeds not because it maximizes PET binding, but because it provides a more favorable balance between modest substrate engagement and enhanced stability in solution.

Such is the nuance of this system; a fine balance between enzyme stability and productive substrate interaction drives PET hydrolysis behavior. The role binding plays for a soluble enzyme with a highly insoluble substrate like PET has previously been conceptualized to abide by the Sabatier principle^32^ stating intermediate strength binding interactions will produce maximal catalytic turnover.^33,34^ We have previously applied this idea for these same supercharged CBMs appended to the cellulase Cel5A and observed strong pH dependent shifts in activity related to surface charge abiding by this design principle.^31^ This same idea was the key driver for the initial hypothesis, but pH sweeps identified no shift in optimal hydrolysis pH and no strong Sabatier-like behavior. Instead, activity was constrained by the optimal pH (6.0) of this Cutinase domain, the same pH where the maximum melting temperature was recorded. This likely points to the thermostability mechanism that led to failure of the original binding hypothesis; although binding can be modulated by net charge, assuming this will track to better catalytic performance in every system is an over-simplification. We have identified a greater nuance for this particular PET hydrolase; enzyme stability is the dominant limiting factor in PET degradation by *Tf.* Cutinase, with binding playing a more subtle secondary role.

Overall, this work demonstrates that charge engineering of appended accessory domains like carbohydrate binding modules can substantially alter the behavior of PET hydrolase fusion enzymes. These appended units not only serve to modulate substrate interactions but can also alter enzyme stability and catalytic persistence through unexpected effects on the ensemble fusion behavior. Although enhanced PET adsorption alone was insufficient to improve hydrolytic performance in this system, the identification of the thermostabilized D2 CBM2a – Cutinase fusion highlights the potential of accessory domain engineering as a strategy to improve polyester hydrolase performance. To harness supercharging effectively for stability engineering, future work such as molecular dynamics simulations are required to understand the interdomain interactions driving stabilization. Future efforts combining highly active PET hydrolases with engineered binding domains and enhanced thermostability may provide a promising route toward more efficient enzymatic PET depolymerization. Collectively, these findings further emphasize that productive PET depolymerization requires balancing substrate interaction with catalytic persistence under conditions favorable for PET hydrolysis, rather than maximizing adsorption alone.

## ACKNOWLEDGEMENTS

This study was supported by Rutgers University, NSF CBET Award (1846797), and USDA Sungrant award (S005193). ADC was supported by the Biotechnology Training Program fellowship that was funded by Rutgers University and the National Institute of General Medical Sciences of the National Institutes of Health under award number T32 GM135141. Original genes used for cloning of CBM fusions were provided by the Department of Energy Joint Genome Institute (DOE-JGI) supported by the Community Science Program Gene Synthesis Award (CSP-503631 Syn). We would like to thank Deanne Sammond (NREL) for help with computational design of supercharged cellulases at the onset of the DOE-JGI award that generated the original CBM designs. Finally, we would like to thank Samantha Shimabukuro (Rutgers University) for lab work support.

