## Supplementary material for "Supercharged binding modules can modulate engineered poly(ethylene terephthalate) hydrolase thermostability and functional persistence": SI Figures and Tables

**Number of SI pages in current PDF (SI-Supplementary Information)**:

**Number of SI figures in current PDF (SI-Supplementary Information):** 14

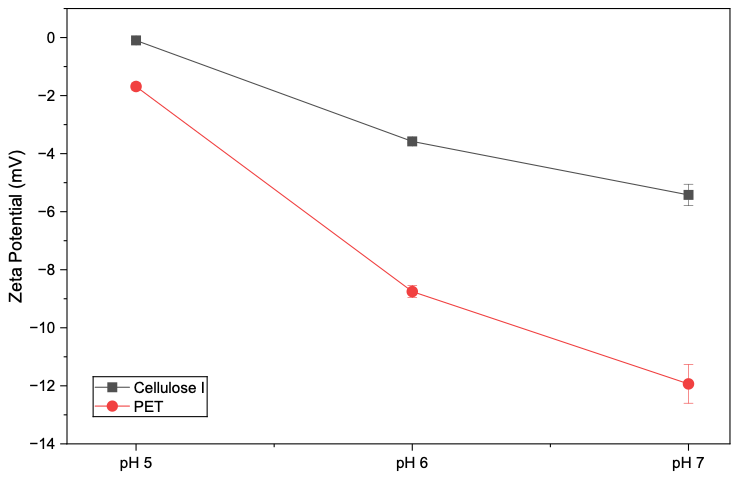

**Supplemental Figure S1. Zeta Potential measurements for crystalline Cellulose – I (Avicel PH-101) and milled PET.** Both powdered substrates were passed through a 50-micron sieve to obtain an equal particle size distribution. Sieved powders were resuspended in 100 mM sodium acetate (pH 5.0) or phosphate (pH 6 or pH 7) buffer at a slurry concentration of 0.5 mg/mL. Zeta potential was measured for quadruplicate samples in a Malvern Zetasizer Nano 25-900 with refractive index values of 1.5 and 1.8 for cellulose and PET respectively. Data points represent the average of four trials, and error bars represent the standard deviation from the mean.

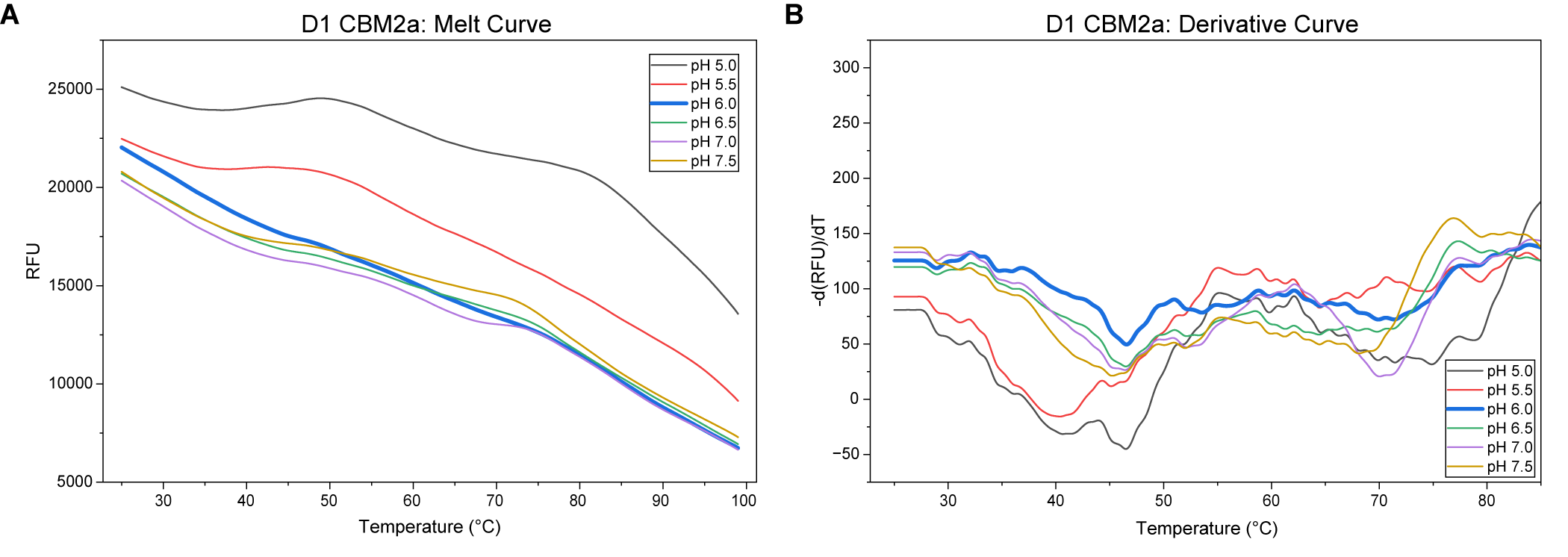
**Supplemental Figure S2. Melt Curve and Derivative Curve for D1 CBM2a – Cutinase**. DSF and subsequent data analysis was performed in a manner consistent with the description listed in the main text. Instability and aggregation artifacts for the D1 CBM2a fusion prevent identification of clear melting trend or melt temperature.

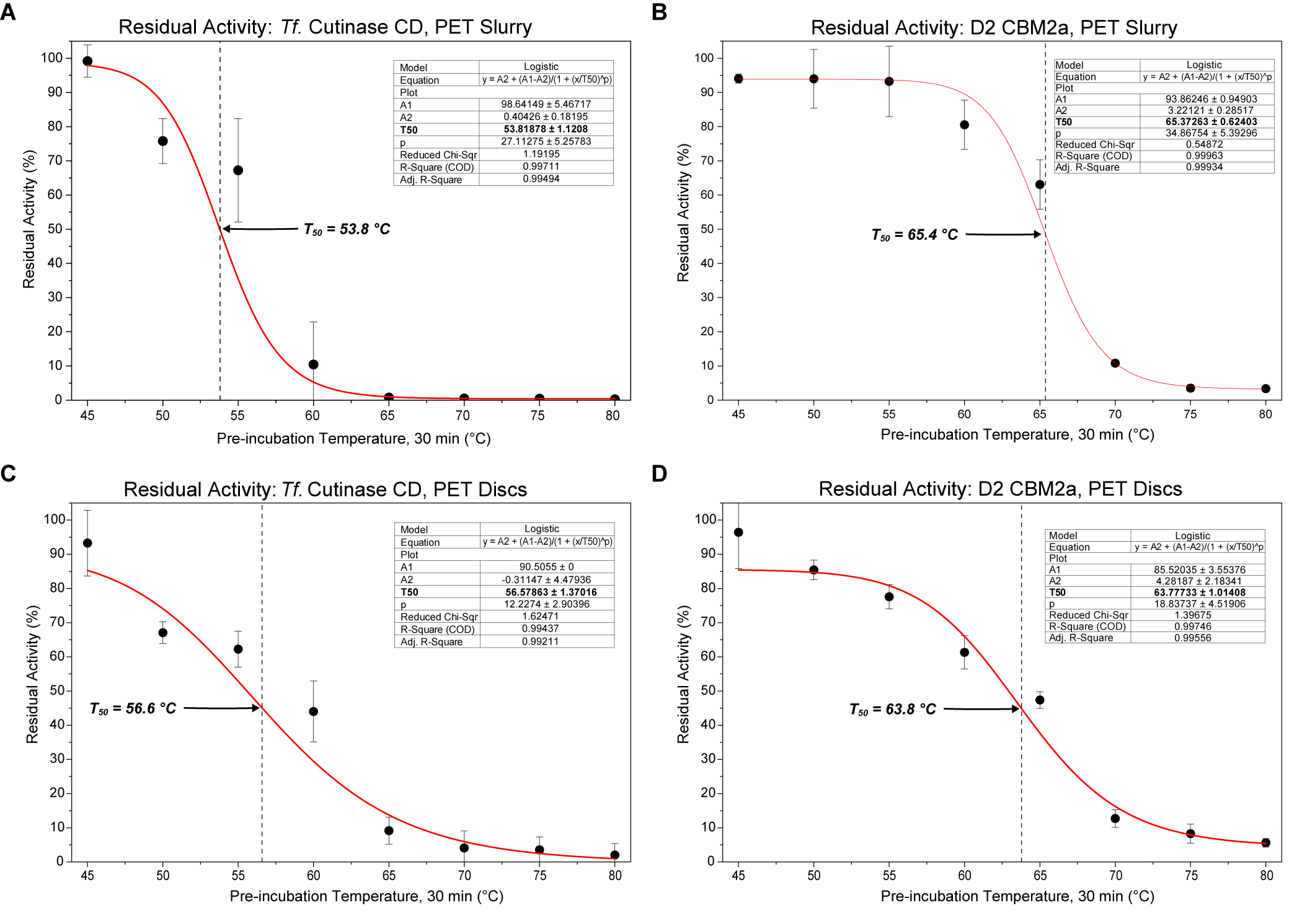

**Supplemental Figure S3.** Residual activity and thermal inactivation behavior of the isolated T. fusca Cutinase catalytic domain (CD) and thermostabilized D2 CBM2a fusion following thermal pre-incubation. Four – parameter logistic curve fits for the isolated Cutinase CD (A) and D2 CBM2a fusion (B) on milled PET and the isolated CD (C) and D2 CBM2a fusion (D) on amorphous PET discs. Assays were performed as described in the main text. Data shown here is originally represented in **Figure 6** of the main manuscript. 4-PL models were fit using Origin and the corresponding to T_50_ values are represented in **Table T2** of the main text.

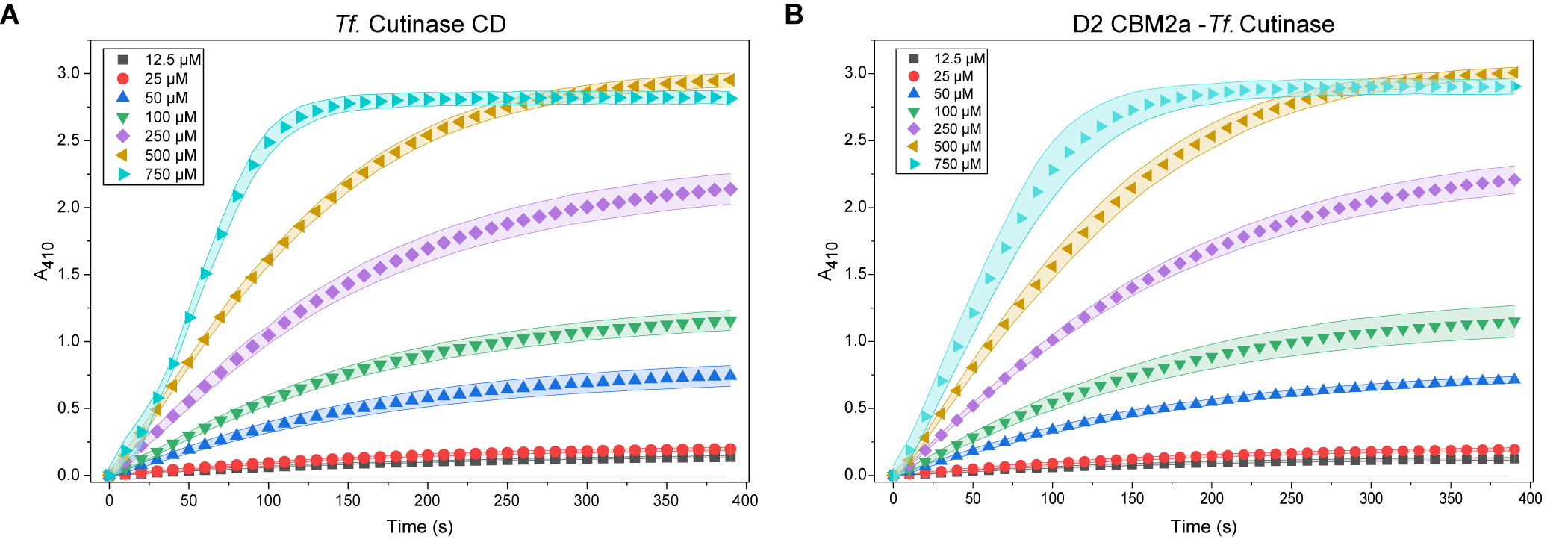

**Supplemental Figure S4. Soluble kinetic characterization of the isolated T. fusca Cutinase catalytic domain (CD) and D2 CBM2a fusion using pNP-acetate.** Reaction progress curves were monitored spectrophotometrically at 410 nm across increasing substrate concentrations. Kinetic traces for the isolated Cutinase CD are shown in (A), while traces for the D2 CBM2a-T. fusca Cutinase fusion are shown in (B). Both enzymes exhibit similar soluble substrate hydrolysis behavior, suggesting that the enhanced PET hydrolysis observed for the D2 fusion does not arise from major alterations in intrinsic catalytic activity on soluble ester substrates.
